# Transgenerational Drought and Methyl Jasmonate Memory Interactively Shape Metabolome and Physiology in Clonal Grass

**DOI:** 10.1101/2025.08.03.668340

**Authors:** Tarun Bhatt, Nikita Rathore, Jaroslav Semerad, Tomas Cajthaml, Dinesh Thakur, Zuzana Münzbergová

**Affiliations:** Department of Botany, Faculty of Sciences, Charles University, Benatska 2, 12800, Prague, Czech Republic, 12800; Institute of Botany, Czech Academy of Sciences, Zamek 1,25243, Pruhonice, Czech Republic; Institute of Microbiology of the Czech Academy of Sciences, Videnska 1083, 142 20 Prague, Czech Republic

**Keywords:** Metabolomics, methyl jasmonate, MeJA memory, drought resilience, clonal plants, transgenerational effects, phenotypic plasticity, drought memory, *Festuca rubra*

## Abstract

As droughts become more common due to climate change, survival may rely not only on a plants immediate response but also on what it has learned from past challenges. However, we still know little about how plants integrate different types of experiences, such as recurrent drought and hormonal cues, from previous generations.

In this study, we examined whether clonal offspring of Festuca rubra plants previously exposed to drought or the stress hormone methyl jasmonate (MeJA) inherited biological memories that help them tolerate new drought stress. Using a factorial design, we combined untargeted LC– MS metabolomics with morpho-physiological measurements to evaluate these memory effects.

We found that each type of memory changed plant metabolism and physiology, but the most notable changes occurred when both memories were present, and plants faced recurrent drought conditions again. This interaction between drought memory, MeJA memory, and current stress did not just add effects, it created entirely new metabolic responses, not seen in any single treatment. These combined memories fine-tuned water conservation, photosynthesis, and extensive metabolomic reshuffling, revealing a deeper level of drought resilience.

Our results uncover a layered memory system in plants where past stresses do not act in isolation but interact to reshape future responses. This offers new insight into how plants prepare for stress and suggests practical strategies for priming drought tolerance across plant generations.

## 1 Introduction

Recurrent droughts are becoming increasingly common under climate change (IPCC, 2023). Unlike isolated drought events, recurrent drought imposes cumulative stress on ecosystems, leading to persistent disruptions in plant functioning and long-term ecological shifts (Jiao et al., 2023a). While recurrent drought has been studied at broader ecosystem and community levels(Hoover et al., 2021; Jiao et al., 2023b; Möhl et al., 2023), but its transgenerational effects remain poorly understood, particularly in clonal grasses. Clonal grasses dominate many temperate and arid ecosystems, and their unique reproductive strategies make them both vulnerable to and potentially resilient against prolonged environmental stress. Understanding how recurrent drought influences not only the immediate generation but also subsequent clonal generations is critical for predicting vegetation dynamics, ecosystem resilience, and carbon cycling under a changing climate.

Grasslands are disproportionately affected by changing drought regimes due to their dependence on seasonal rainfall. Many of these ecosystems are dominated by clonal grasses, whose persistence and productivity are tightly linked to water availability(Luo et al., 2023; Lv et al., 2024). Clonal reproduction allows these species to maintain population continuity during stress events but may also transmit physiological and developmental legacies across generations. As recurrent drought becomes more frequent, it is crucial to understand how these grasses respond not just within a single generation but over subsequent vegetative cycles. Such transgenerational responses could fundamentally reshape grassland composition, resilience, and function. Yet, how recurrent drought translates into metabolomic and morphophysiological reprogramming in clonal grasses remains unclear, limiting our ability to link ecosystem-level patterns to plant-level processes.

Phytohormones play a central role in orchestrating plant responses to drought stress. Abscisic acid (ABA), salicylic acid (SA), jasmonic acid (JA), and ethylene regulate processes such as stomatal closure, root elongation, oxidative stress mitigation, and systemic defense signaling (Samanta et al., 2024). Under recurrent drought, hormonal pathways often exhibit heightened responsiveness, reflecting prior stress exposure (Avramova, 2019a). Among these hormones, methyl jasmonate (MeJA) is of particular interest because it enhances drought tolerance by modulating root and shoot growth, improving water-use efficiency, and activating antioxidant defenses (Mohamed & Latif, 2017). It also regulates key developmental processes such as senescence, highlighting its broad role in coordinating stress adaptation (Hewedy et al., 2023a). This makes MeJA a strong candidate for inducing stress memory and potentially transmitting adaptive advantages across generations. Despite this importance, the role of MeJA in shaping transgenerational drought memory remains poorly resolved, particularly in perennial, clonally reproducing grasses.

Evidence increasingly suggests that plants are not passive responders to stress but can “remember” prior exposures through the establishment of hormonal memory. Repeated drought events leave behind a physiological and molecular imprint that enables plants to mount faster and stronger defenses when stress reoccurs(Kambona et al., 2023a; Liu et al., 2022). Despite this progress, one critical question remains unanswered can methyl jasmonate (MeJA), a key regulator of both drought tolerance and developmental processes, itself establish such a drought-related memory. Even more intriguingly, in clonal plants, could this memory interact with drought memory. The interaction between recurrent drought and hormone memory is expected to be more than just an additive response it is a dynamic feedback system that equips plants with an enhanced capacity for survival under recurring stress.

Another critical aspect that remains unexplored in the context of drought and MeJA memory in clonal plants is how whole-plant metabolomic profiles shift and how these align with morpho-physiological responses. Whole metabolomics is powerful in this context because it captures the entire biochemical reprogramming triggered by recurrent drought and MeJA, including not only hormones but also osmolytes, antioxidants, and secondary metabolites. Unlike targeted assays, it offers an unbiased and integrative perspective, revealing subtle shifts that may precede visible morpho-physiological changes. Comparing metabolomic diversity and composition with plant performance allows us to test whether biochemical memory predicts resilience, thereby bridging cellular biochemicals with ecological outcomes.

We hypothesize the following: **H1** – Exposure to recurrent drought (D2) strongly influences the metabolomic and morpho-physiological traits of *F. rubra*, representing immediate stress-induced changes. **H2**-Prior exposure to recurrent drought (D1) and methyl jasmonate (MeJA, M1) induces transgenerational memory in clonal offspring, leading to persistent reprogramming of the metabolome and modulation of key morpho-physiological traits such as specific leaf area (SLA), relative water content (RWC), and chlorophyll dynamics thereby enhancing plant responses to recurrent drought stress even in the absence of current drought. **H3** – Recurrent drought and methyl jasmonate memories will interact with current recurrent drought such that combinations of past and present stress exposures will produce synergistic or compensatory effects on plant metabolome and morpho-physiology. **H4**-Finally, we hypothesize that metabolomic profiles and morphophysiological traits are tightly linked, reflecting coordinated memory-driven responses across molecular and phenotypic levels that enhance drought resilience.

## 2. Materials and Methods

### 2.1 Study system

We used ramets of *F. rubra* from our previous experiment (Bhatt et al., n.d.-a) for this study. The previous experiment involved a factorial combination of drought (yes/no) and MeJA (yes/no) treatments and 10 replicates per treatment. *F. rubra* ramets were grown in pots with natural grassland soil under controlled conditions. Drought stress was applied in two phases, with recovery phases in between, while MeJA was applied to the soil during drought phases and not during the recovery phases. Control plants received regular watering. The MeJA soil application was chosen to ensure consistent distribution and minimize environmental loss. More details from the previous experiment are given in (Text S1).

In this experiment, ramets previously subjected to different drought and MeJA treatments (referred to as parental treatment) were grown in pots containing soil exposed to either drought or MeJA + drought in the previous experiment. Six replicates of ramets from the four parental treatments were grown in each soil with different memories (i.e., 2 parental drought treatments × 2 parental MeJA treatments × 2 soil memories × 2 current drought × 6 replicates). Although soil memory (SM) was included in the experimental design to account for potential legacy effects from the previous generation, it was not a primary focus of this study. SM showed statistically significant effects only in the metabolomic composition analyses and was not significant in any other aspect of the study. Consequently, SM is not presented in the main text, and its results are presented only in (Table S1). We focus only on intrinsic plant memory mechanisms, specifically recurrent drought (D2), drought memory (D1), and MeJA memory (M1).

Three of these replicates were well watered, and the remaining three were subjected to recurrent drought cycles followed by recovery phases. This included 30 day cycles of drought and recovery (Drought stage 1 - Recovery stage 1 - Drought stage 2 - Recovery stage 2). No MeJA treatment was applied, as the focus was on investigating the legacy effects of prior MeJA treatment on the metabolome of next-generation ramets under recurrent drought conditions.

The plants were grown in 200 ml pots. One *F. rubra* ramet was planted into each pot. All planted ramets were of similar age and size (3 weeks old, 3.5-4 cm tall). They were grown under long-day conditions (16 hours of daylight) in a growth chamber (17–20 °C Day and 7– 10 °C Night), reflecting peak growing season conditions at the original locality. The plants were allowed to establish for 21 days before starting the experiment.

### 2.2 Plant performance

Plant physiology was characterized by chlorophyll content, chlorophyll fluorescence (Fv/Fm), specific leaf area (SLA), leaf dry matter content (LDMC), and leaf relative water content (RWC). These traits provide critical information about how plants structurally and physiologically adapt to drought stress(Maxwell & Johnson, 2000; Perez-Harguindeguy et al., 2013a).

Chlorophyll content and Chlorophyll fluorescence (Fv/Fm) were measured at the end of each stress and recovery phase. SLA, LDMC, and RWC were measured only once, at the end of the second recovery phase, due to their destructive nature, which prevented repeated measurements.

#### 2.2.1 Chlorophyll Content and Chlorophyll Fluorescence

Chlorophyll is an essential pigment crucial for photosynthesis, and its detection in plants is a significant indicator of their chlorophyll fluorescence and overall well-being (Muller et al., 2011). In our study, we utilized a chlorophyll CCM 300 meter to quantify the chlorophyll content in plant leaves. Three randomly selected leaves were chosen from each pot, and the chlorophyll content was measured according to the instructions provided by the manufacturer once per leaf. The mean of these 3 values has been used as the dependent variable in the subsequent analyses.

Chlorophyll degradation can decrease the maximum efficiency of Photosystem II (PSII) known as the Fv/Fm ratios, decreasing the net photosynthetic rate (Chungloo et al., 2023). Photon System Instrument FluorPen FP-100 MAX determined the chlorophyll fluorescence of PSII. Healthy plants adapted to darkness have chlorophyll fluorescence ratios ranging from 0.75 to 0.85 (Björkman & Demmig, 1987). Lower values indicate damage to the system. chlorophyll fluorescence was measured in the afternoon, and a 1-hour dark acclimation was achieved by using the manufacturer provided clips. Chlorophyll content and chlorophyll fluorescence were measured across four key phases: stress phase 1, recovery phase 1, stress phase 2, and recovery phase 2. Based on the correlation analysis (Figure S1), two representative variables were selected to reduce redundancy and multicollinearity: CCS2 (Chlorophyll Content during Stress Phase 2) and CFR1 (Chlorophyll Fluorescence during Recovery Phase 1).

CCS2 showed strong correlations (r ≥ 0.7) with other chlorophyll content traits from stress phase 2 and both recovery phases, as well as with chlorophyll fluorescence traits from both stress phases. This suggests that CCS2 effectively captures broader chlorophyll related variation across phases. Similarly, CFR1 was highly correlated with CFR2 (Chlorophyll Fluorescence during Recovery Phase 2), making it a suitable representative of the recovery-phase fluorescence response.

#### 2.2.2 SLA, LDMC, and RWC

SLA (Specific leaf area) and LDMC (Leaf dry matter content) are frequently used to predict drought resistance because they essentially represent a trade-off between conservation and rapid resource uptake (Blumenthal et al., 2020; Perez-Harguindeguy et al., 2013b). RWC (Relative water content) serves as the most appropriate measure for assessing a plant’s water status, providing insight into cellular water deficit and the impact of Osmotic Adjustment on maintaining cellular hydration during drought stress (Barrs & Weatherley, 1962).

Due to the extremely narrow and naturally folded morphology of *F. rubra* leaves, accurately determining leaf area from scanned images proved unreliable. To standardize measurements, we selected approximately 4 cm-long segments from individual leaves per pot. Fresh weight was recorded for each segment. Because the leaves could not be fully unfolded without damage, we measured both the length and width of each segment using a magnifying glass. Leaf area was then estimated using the formula: length × width × 2, accounting for the folding of the leaf blade. Subsequently, the leaf segment was placed in a Petri plate filled with water overnight in a refrigerator to rehydrate. After the leaves had been rehydrated, any extra water on the surface was carefully removed, and the leaves were weighed to determine their turgid weight. The leaves were then dried in a 65°C oven for two days to determine their dry weight. Using these measurements, we calculated the SLA by determining the ratio of leaf area (mm^2^) to dry mass (mg), LDMC by dividing the dry leaf mass (mg) by the turgid mass (g), and Relative water content RWC as RWC (%) = [(Fresh weight - Dry weight) / (Turgid weight – Dry weight)] × 100.

#### 2.2.3 Metabolite Profiling

##### 2.2.3.1 Plant sampling

Samples were collected at the end of the second drought phase. Approximately 20 leaves per individual from multiple ramets per individual were sampled and pooled. Leaves were carefully excised using sterile scissors, then wrapped in aluminium foil and immediately placed in liquid nitrogen for snap freezing and temporary storage. After sampling all the plants, the samples stored in liquid nitrogen were transferred to a -80°C freezer for storage until further analyses.

##### 2.2.3.2 Methanol extraction

We took ∼100 mg of each sample. We placed it into 1.5 ml microcentrifuge tubes (MCT) containing 3 metal beads and 500 µL of an extraction solvent composed of methanol and water (80:20). The samples were homogenized using a homogenizer. After homogenization, samples were thoroughly vortexed for 10 seconds, followed by a 24-hour incubation at -20 °C. After incubation, the tubes were vortexed again and centrifuged at 13000 rpm for 15 minutes. The supernatant was carefully transferred into new tubes. The same steps of the extraction process, from adding the solvent, were repeated once to ensure complete metabolite extraction. The supernatants from both extraction rounds were pooled together. From this combined pool, 320 µL was transferred to a new tube, and 80 µL of a water/formic acid solution (99.9/0.1 v/v, respectively) was added to the tube. The samples were then stored at -20 °C for at least 48 hours. Three blanks were processed alongside the samples to detect solvent-related contamination, following all extraction steps but excluding leaf tissue. Additionally, quality controls were created by pooling 20 µL from each sample extract.

##### 2.2.3.3 Liquid Chromatography Data Acquisition

Chromatographic separation was performed at 40 °C using a 6546 LC/Q-TOF (Agilent) system equipped with an Acclaim® RSLC 120 column (150 × 2.1 mm, 2.2 μm particle size; Thermo Fisher Scientific). The chromatographic run began with an initial mobile-phase composition of 95 % A, maintained for the first 3 minutes. From 0–3 minutes, the proportion of A was linearly reduced to 82.7 %, then decreased to 76 % by the 10-minute mark. A steep linear gradient followed between 10 and 17 minutes, dropping sharply from 76 % to 5 %, which was held constant for one minute. A rapid re-equilibration occurred between 18 and 18.1 minutes, raising A back to 95 %, and this composition was maintained through 20 minutes. The entire sequence was executed for both positive and negative ionization modes. MS/MS analysis was also performed in the same settings in a data-dependent mode, with 4 analyses conducted from QC samples.

##### 2.3.3.4 Spectral processing

Spectral processing of the LC-MS data was performed to extract relevant features for metabolite identification and quantification. The raw data, which contained metabolites and their concentrations across multiple time points, was processed using the MetaboAnalystR Package of R (Pang et al., 2024). Several preprocessing steps were further applied to ensure the quality and reliability of the acquired data. First, data were filtered to remove features with more than 85% missing values, which helps eliminate unreliable or incomplete data. Missing values were then imputed using the 1/5^th^ of the lowest value (Pang et al., 2024). This step is essential to minimize the impact of missing values, which may arise due to biological variability or technical limitations in detection sensitivity. Subsequently, the data were quantile normalized, log transformed and scaled to stabilize variance and ensure comparable distributions across samples. However, for estimating metabolite diversity metrics (richness, Shannon diversity, and evenness), the processed data were deemed unsuitable due to transformations that alter the original distribution of metabolite presence. Therefore, diversity estimates were calculated using the unprocessed data to preserve the integrity of presence-absence and abundance patterns.

### 2.3 Data analysis

All the metabolome analyses have been done separately for positive and negative ionization modes as in previous studies (e.g. Calderón-Santiago et al., 2016; Nordström et al., 2008; Tian et al., 2013).

#### 2.3.1 Metabolite diversity and Uniqueness

To evaluate the effects of transgenerational memory of drought and Methyl Jasmonate (MeJA) on metabolite diversity, we analyzed three key diversity indices, i.e., metabolite richness, Shannon diversity, and Hill evenness (HillEven). Richness was estimated as the total number of detected metabolites in each of the samples. Shannon diversity and HillEven were estimated using the Chemodiv Package of R (Petrén et al., 2023). These indices were chosen because they provide complementary insights into the metabolome: Shannon diversity captures both the number and relative abundance of metabolites, reflecting metabolic complexity, while Hill evenness quantifies how evenly metabolite abundances are distributed, which is critical for understanding dominance patterns in untargeted metabolomics datasets. Additionally, metabolomic uniqueness was evaluated by calculating the average Euclidean distance of each sample from all others based on standardized metabolite intensities (Ren et al., 2015) .

Linear models were fitted using a full factorial design incorporating all treatments: D1, D2, M1, and their two-way interactions. Before analyses, Richness, Shannon Diversity, and Hill Evenness were log-transformed to fulfil the data normality assumption. Because the results of Hill evenness largely matched the results of Shannon diversity in our analyses (r = 0.96), they are not reported further. They are only shown in supporting information (Table S2 and Figure S2, S3).

#### 2.3.2 Multivariate analyses

Multivariate analyses were applied to examine the interactive effects of treatments and morpho-physiological traits on the metabolomic composition. Redundancy Analysis (RDA) as implemented in the vegan package in R was used throughout to explore these relationships, using standardized data for all dependent variables (i.e., metabolites). A full model including all main effects of D1, D2, M1, and their interactions was constructed. The significance of each predictor was determined using permutation tests (n = 999), and variance partitioning was used to quantify their contribution to overall variation. A similar RDA approach was used to investigate how morpho-physiological traits influence metabolomic composition and how the treatments affected plant morpho-physiological traits.

#### 2.3.3 Effect of treatments on each metabolite

To visualize the overlap and specificity of metabolite responses across treatment groups, we quantified the number of significantly affected metabolites (FDR-adjusted *p* < 0.05) for each main effect (D1, M1, D2) and their pairwise interactions (D1×D2, D1×M1, D2×M1). Significant metabolites were extracted from the linear model output, and treatment-specific sets of metabolites were defined. A binary presence/absence matrix was constructed across all groups, enabling classification of each metabolite as either unique (significant in one group only) or shared (significant in multiple groups). The resulting counts were visualized using bar plots in R (version 4.2.1). To visualize changes in metabolite content, a heatmap was constructed based on group-level averages of significant metabolites, with hierarchical clustering. Also, due to the huge number of significant metabolites, only the top 100 were used in the heatmap.

#### 2.3.4 Annotation of significantly affected metabolites

We annotated the metabolites significantly affected by climate. For this, data-dependent LC-MS/MS analysis was performed on a pooled sample (used as a quality control in LC-MS analysis) to collect the necessary data. Subsequently, we used MS-DIAL (Tsugawa et al., 2015) software to putatively annotate the metabolites detected in LC-MS/MS mode data. LC-MS/MS data were processed by aligning retention times, detecting peaks, and deconvoluting spectra. The annotation was then done by matching detected compounds against freely available libraries at https://systemsomicslab.github.io/compms/msdial/main.html. A mass accuracy threshold of 10 ppm was used in the annotation process. After annotation, the metabolites were assigned to their chemical classes.

## 3 Results

### 3.1 Metabolite Diversity

The effect of recurrent drought (D2) and its interaction with drought memory (D1) was statistically significant (Table 1) in the positive ionization mode. Metabolite richness declined under D2, indicating a narrowing of the metabolomic profile in response to stress. The significant interaction between D1 and D2 suggests that the benefits of drought memory on metabolomic diversification are conditional upon subsequent drought exposure. Specifically, drought memory (D1) increased metabolite richness under non-stress conditions, but had the opposite effect under current drought, where richness was lowest in plants that had been previously exposed to drought (Figure 1a).

**Figure 1:**
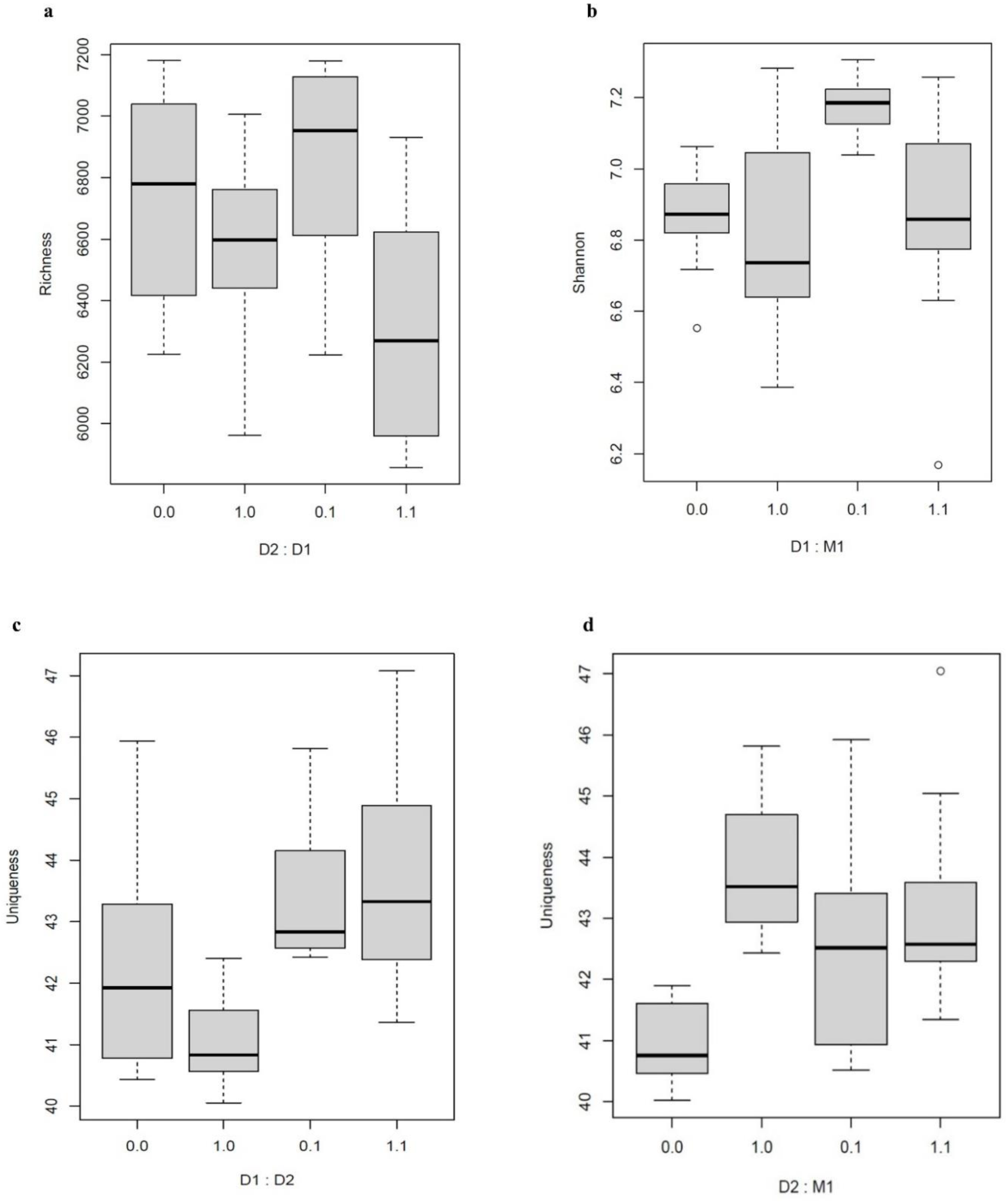
Boxplots showing the effects of interactions of recurrent drought (D2), drought memory (D1), and MeJA memory (M1) in positive mode on metabolome (a) Richness, (b) Shannon diversity, and (c, d) Uniqueness in *F. rubra*. The X axis shows the levels of the interacting treatment, with 0 indicating absent and 1 present treatment (recurrent drought, drought memory or MeJa).

**Table 1:** Effects of D1, D2, M1, and their interactions on Richness, Shannon diversity, and Uniqueness in both positive and negative modes. The variation explained is shown for the significant predictors (p < 0.05).

| Treatment | Richness |  |  |  | Shannon |  |  |  | Uniqueness |  |  |  |
| --- | --- | --- | --- | --- | --- | --- | --- | --- | --- | --- | --- | --- |
|  | Positive mode |  | Negative mode |  | Positive mode |  | Negative mode |  | Positive mode |  | Negative mode |  |
|  | P-value | Variation explained | P-value | Variation explained | P-value | Variation explained | P-value | Variation explained | P-value | Variation explained | P-value | Variation explained |
| D1 | 0.43 | - | 0.7 | - | 0.011 | 12.29 | 0.83 | - | 0.20 | - | 0.31 | - |
| D2 | <0.001 | 23.6 | 0.54 | - | 0.927 | - | 0.008 | 14.41 | 0.001 | 28.72 | 0.001 | 44.07 |
| M1 | 0.146 | - | 0.39 | - | 0.009 | 12.82 | 0.24 | - | 0.25 | - | 0.094 | - |
| D1×D2 | 0.038 | 6.59 | 0.001 | 21.34 | 0.343 | - | 0.14 | - | 0.023 | 5.44 | 0.014 | 5.88 |
| D1×M1 | 0.659 | - | 0.85 | - | 0.049 | 6.63 | 0.36 | - | 0.094 | - | 0.097 | - |
| D2×M1 | 0.77 | - | 0.7 | - | 0.528 | - | 0.9 | - | 0.001 | 11.93 | 0.02 | 4.78 |
| D1×D2×M1 | 0.309 | - | 0.484 | - | 0.38 | - | 0.954 | - | 0.074 | - | 0.012 | 5.39 |

For Shannon diversity, both D1 and M1 had statistically significant effects in the positive ionization mode (Table 1), indicating that each memory treatment independently affects metabolite diversity. D1 individually caused a decline in Shannon diversity, while M1 caused an increase. Their interaction (D1 × M1) was also significant (Table 1), indicating that the effect of M1 on Shannon diversity depends on D1. Specifically, M1 caused an increase in Shannon diversity in the absence of D1 memory, but the combined presence of D1 and M1 memories shaped Shannon diversity by negating the effects of each other and bringing the Shannon diversity to control levels (both D1 and M1 are absent) (Figure 1b). In contrast, D2 did not significantly influence Shannon diversity in the positive mode but showed a strong effect in the negative ionization mode (Table 1), where it significantly decreased Shannon diversity (Figure S4b), reflecting a more uneven distribution of metabolites under stress.

### 3.2 Metabolomic Uniqueness

The recurrent drought (D2) was a significant predictor of metabolomic uniqueness in both positive and negative ionization modes (Table 1), while M1 and D1 were not. A significant interaction between D1 and D2 was also detected (Table 1), showing that metabolite uniqueness declined under D1 alone but increased notably when drought memory plants were exposed to current drought (Figure 1c). Also, the interaction between D2 and M1 was statistically significant (Table 1). Plants without MeJa memory without current drought showed the lowest uniqueness, plants without MeJa memory with current drought showed the highest uniqueness and plants with MeJa memory, independent of current drought being in between (Figure 1d). While the figures show results for the positive mode only, the results for the negative mode are largely similar (Figure S4c & S4d)

The triple interaction (D1 × D2 × M1) was significant for metabolic uniqueness in negative mode. Plants with both drought and MeJa memory grown under drought exhibited the highest metabolic uniqueness compared to all the other treatments (Figure S5).

### 3.3 Determinants of metabolome composition

D2 was the strongest predictor of metabolome composition in both positive and negative ionization modes (Table 2). M1 and D1 (Table 2) also had significant and independent impacts on metabolite profiles in both modes, though their effects were comparatively lower than D2. The interactions D1×M1 and D2×M1 were also statistically significant (Table 2), indicating that the combination of stress memory with current drought or between both memory types altered metabolite composition. In contrast, the interaction between D1 and D2 did not show a significant effect in either mode.

**Table 2:**
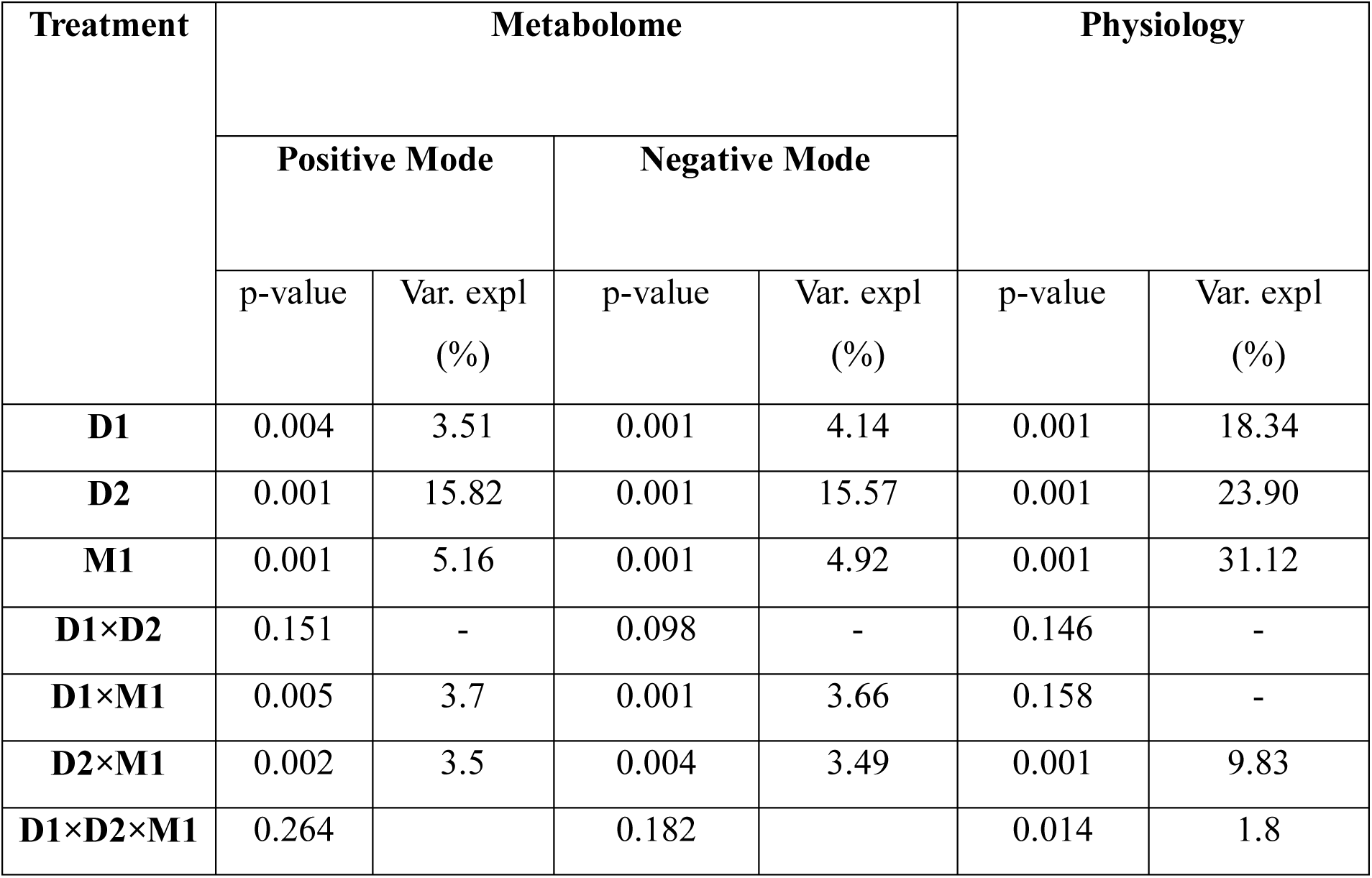
Summary of Redundancy Analysis (RDA) results showing the effects of drought memory (D1), recurrent drought (D2), MeJA memory (M1), and their interactions on metabolomic profiles (in positive and negative ionization modes) and morpho-physiological traits. The table reports the p-values and percentage of variation explained for each significant predictor. Only significant values (p ≤ 0.05) are associated with variation explained (%); dashes (–) indicate non-significance.

| Treatment | Metabolome |  |  |  | Physiology |  |
| --- | --- | --- | --- | --- | --- | --- |
|  | Positive Mode |  | Negative Mode |  |  |  |
|  | p-value | Var. expl (%) | p-value | Var. expl (%) | p-value | Var. expl (%) |
| D1 | 0.004 | 3.51 | 0.001 | 4.14 | 0.001 | 18.34 |
| D2 | 0.001 | 15.82 | 0.001 | 15.57 | 0.001 | 23.90 |
| M1 | 0.001 | 5.16 | 0.001 | 4.92 | 0.001 | 31.12 |
| D1×D2 | 0.151 | - | 0.098 | - | 0.146 | - |
| D1×M1 | 0.005 | 3.7 | 0.001 | 3.66 | 0.158 | - |
| D2×M1 | 0.002 | 3.5 | 0.004 | 3.49 | 0.001 | 9.83 |
| D1×D2×M1 | 0.264 |  | 0.182 |  | 0.014 | 1.8 |

### 3.4 Effect of MeJA memory and its interaction with drought on specific metabolites

In the positive ionization mode, D2 induced the most significant metabolic changes, uniquely affecting 475 metabolites and sharing 935 with other treatments, underscoring its central role in metabolomic reprogramming (Figure S7a). M1 had the second largest impact, with 90 unique metabolites and 389 shared, while D1 had the smallest individual effect with 19 unique, 249 shared metabolites. Interaction terms also demonstrated pronounced metabolic responses. The D2 × M1 interaction influenced the most metabolites among combinations (39 unique, 205 shared), followed by D1 × M1 (19 unique, 192 shared), while D1 × D2 showed minimal effect (0 unique, 1 shared).

In the negative ionization mode (Figure S7b), D2 again triggered the strongest metabolic shifts (462 unique, 952 shared), with M1 (73 unique, 396 shared) and D1 (30 unique, 311 shared) trailing behind. Similarly, interactions revealed substantial effects: D2 × M1 (54 unique, 170 shared), D1 × M1 (28 unique, 245 shared), and again, D1 × D2 (0 unique, 1 shared).

These findings highlight D2 as the primary driver of metabolic reprogramming, while memory effects from M1 and D1 also shape distinct biochemical landscapes. The notably high number of unique metabolites in D2 × M1 and D1 × M1 interactions indicates a non-additive, context-dependent integration of stress memory with ongoing drought stress.

The heatmap of the top 200 differentially accumulated under different conditions and annotated metabolites in both positive and negative mode (Figure 3). Clusters 1-4 in the positive ion mode heatmap (Figure S8a,b,c & d) illustrate distinct metabolic strategies shaped by MeJA memory in *F. rubra*. Cluster 1 metabolites, including phenolic compounds, flavonoids, curcuminoids, saccharolipids, and linear diarylheptanoids, are upregulated in M1D2-treated plants, revealing a jasmonic acid (JA) specific metabolic imprint, while D2, D1D2, and M1D1D2 treatments show downregulation, suggesting that drought memory alone or in combination interferes with this JA-associated response. Cluster 2 metabolites are selectively upregulated in both M1 and M1D2, including alkaloids, terpenoids, isoflavonoids, lignans, and oligopeptides, indicating that this set is a robust MeJA memory marker that does not require current stress for activation. This exclusive response in MeJA primed plants makes Cluster 2 a clear biomarker for JA-driven metabolic readiness. In contrast, Cluster 3 metabolites such as phosphocholines, salicylamides, diacylglycerols, ether lipids, and flavonoid derivatives are selectively downregulated only in M1 and M1D2, whereas all other treatments, regardless of memory, show upregulation. This suggests that MeJA priming deliberately suppresses this non-primed, general stress response pathway, likely to conserve energy and direct metabolism toward more specialized defenses. Lastly, Cluster 4 is uniquely downregulated in M1D2, whereas D2, D1D2, and M1D1D2 exhibit strong upregulation of their lipid and redox related metabolites including ceramides (Cer_NDS, CerP), fatty alcohol esters, gamma amino acids, and pyridoxines. This pattern indicates that MeJA memory strategically suppresses broad stress induced metabolic activation in favor of targeted defense, reinforcing its role in refining and focusing the plant’s metabolic response under recurring drought conditions.

In negative ion mode (Figure 3b, also Figure S8e&f), Cluster 1 is selectively upregulated in M1, and M1D2 includes metabolites such as flavonoids, coumarins, limonoids, glycerophospholipids, oligosaccharides, and rotanones, establishing it as a MeJA memory-specific metabolic signature. In contrast, D2, D1D2, and M1D1D2 show clear suppression of this cluster, indicating that drought memory alone or even in combination with MeJA fails to activate, and may actively repress, these pathways. This underscores the specificity and strength of MeJA driven priming in shaping targeted metabolic defense readiness. Conversely, Cluster 2 displays the opposite trend M1D2 exhibits strong downregulation of broad stress response metabolites such as alkaloids, phosphatidic acid, alpha and N-acyl amino acids, methoxyphenols, and dihexosylceramides, while D2, D1D2, and M1D1D2 show robust upregulation. This contrast highlights a MeJA memory induced suppression of diffuse metabolic activation, reinforcing a strategic shift in *F. rubra* toward focused, energy efficient defense responses during drought stress.

### 3.5 Relationship between plant morpho-physiological traits and metabolite profiles

The redundancy analysis (RDA) shows that SLA and RWC were the plant traits having the strongest effects on metabolite profiles under both positive and negative ionization modes (Table 3). Chlorophyll content (CCS2) also contributed to explaining metabolomic variation, but to a lesser extent and LDMC had no significant effect on metabolite patterns.

**Table 3:** Effects on plant traits on metabolic composition for positive and negative ionization modes assessed using redundancy analysis (RDA).

|  | Positive mode |  | Negative mode |  |
| --- | --- | --- | --- | --- |
|  | P | Var. expl (%) | P | Var. expl (%) |
| **SLA** | 0.001 | 12.26 | 0.001 | 11.11 |
| **LDMC** | 0.112 | - | 0.106 | - |
| **RWC** | 0.003 | 4.18 | 0.001 | 4.07 |
| **CCS2** | 0.015 | 3.32 | 0.016 | 2.92 |
| **CFR1** | 0.135 | - | 0.136 | - |

### 3.6 Determinants of plant morpho-physiological traits

The redundancy analysis (RDA) showed that M1 was the strongest predictor of morpho-physiological traits in *F. rubra,* followed by D2 and D1 (Table 2). The interaction between D2 and M1 was also statistically significant, while other interactions (D1×D2 and D1×M1) were not significant. The triple interaction between D1×D2×M1 was also significant (Table 2). This suggests that the way D1 affects dependence on M1 changes depending on whether the plants are experiencing recurrent drought (D2).

LDMC, RWC, and CCS2 increased with M1 and D2×M1, indicating positive associations with these treatments. In contrast, SLA pointed in the opposite direction, suggesting a likely trade-off under those same treatments. CFR1 shows a positive association with M1, as both vectors point in the same leftward direction on the RDA biplot. This suggests higher CFR1 values under M1 treatment (Figure 2).

**Figure 2:**
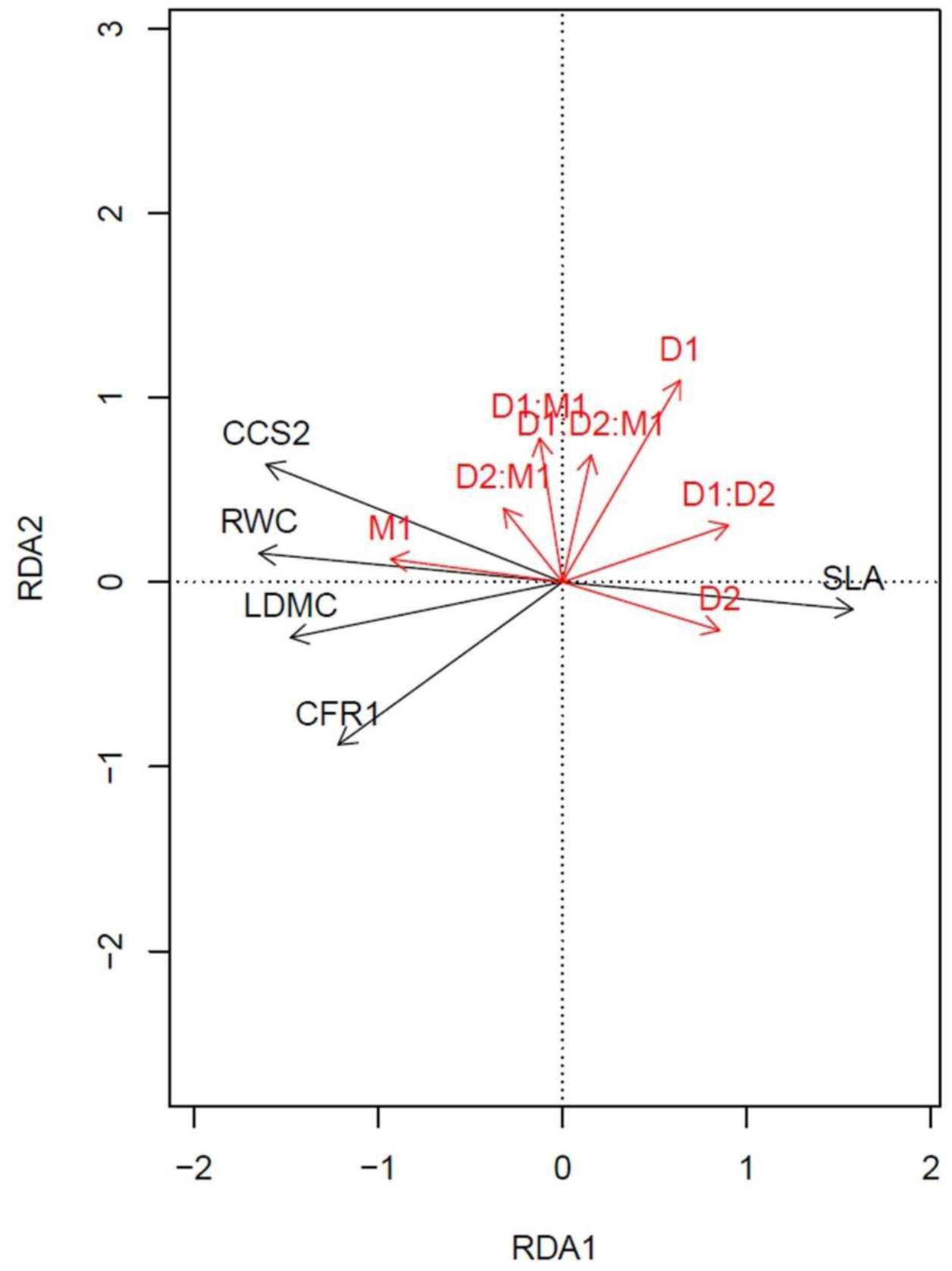
RDA biplot illustrating the relationships between treatment variables (shown as red arrows) and morpho-physiological traits (shown as grey arrows) in *F. rubra*.

**Figure 3:**
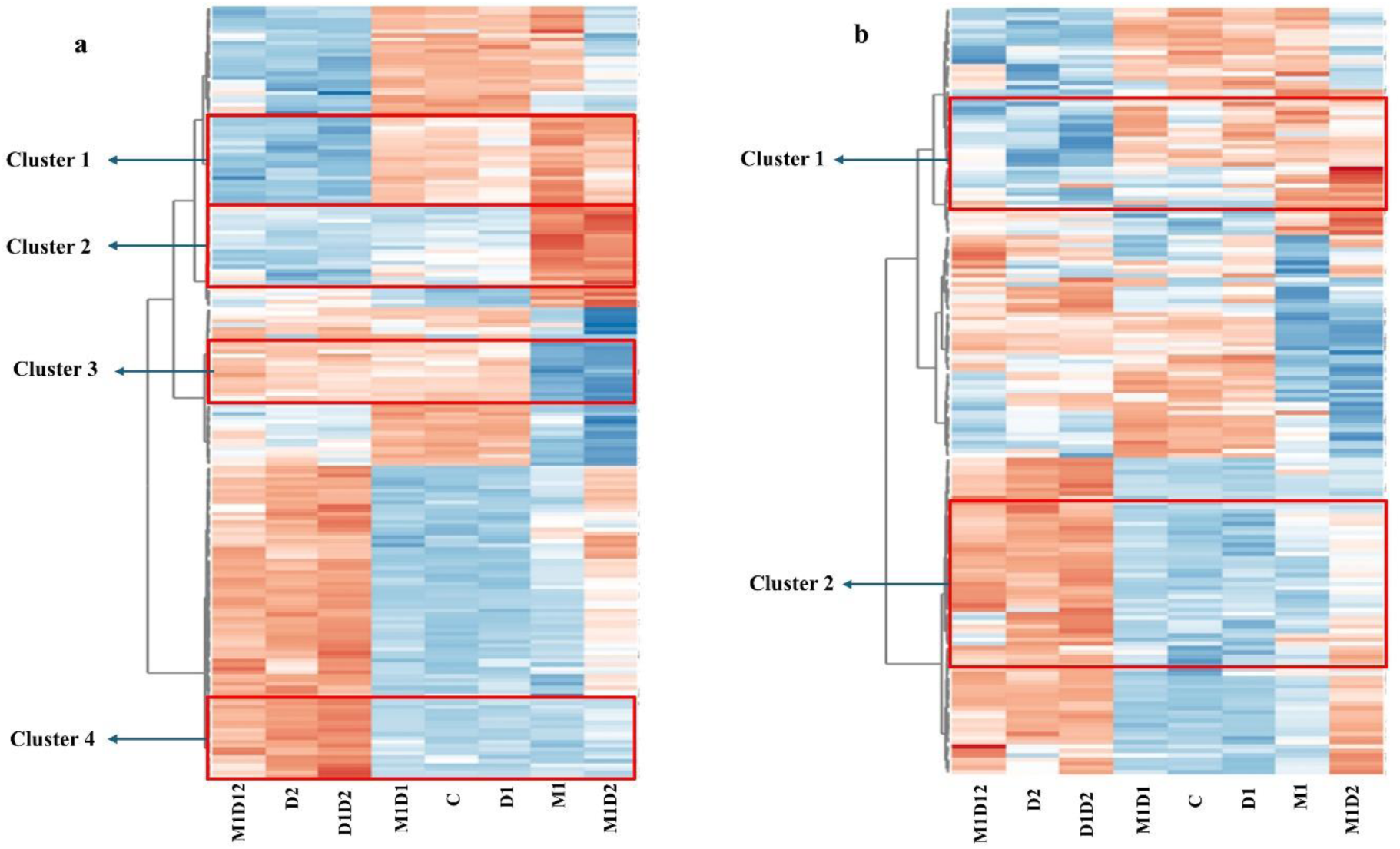
Heatmap displaying the scaled intensity values of differentially accumulated metabolites across various treatment groups in *F. rubra* (M1, D1, D2, i.e. MeJA memory, drought memory and current drought respectively), others are the plants which got combination M1, D2 and D2 treatment, while missing code in combination indicates that plants where not treated by the given treatment. C indicates Control, i.e., no drought or MeJa treatment memory, and also in well-watered conditions in both positive mode (a) and negative mode (b). The clustering was performed using hierarchical clustering based on Euclidean distance and complete linkage. The columns represent plants with different memory and current stress conditions. The orange color in the plot represents a higher accumulation of a metabolite, while the blue color represents a lower accumulation. Also, the zoomed image of marked clusters is provided in the Figure S8 with metabolite names in marked clusters.

## 4. Discussion

This study offers a comprehensive analysis of transgenerational stress memory by integrating metabolomic and morpho-physiological traits in clonal offspring of *Festuca rubra*. We specifically examined the effects of parental drought memory (D1), parental methyl jasmonate (MeJA) priming memory (M1), and their interaction with recurrent drought applied in the second generation (D2). Our results reveal that these stress legacies act both independently and interactively to reprogram metabolic diversity and structural traits. A key novelty lies in showing that memory effects are strongly interaction-dependent, producing conditional shifts shaped by environmental context. Importantly, MeJA memory stabilized drought-induced alterations, acting as a modulator of systemic resilience. By linking metabolomic reprogramming with coordinated morphological responses, our study advances understanding of transgenerational memory and highlights metabolomic memory as a heritable trait in clonal plants.

### 4.1 Recurrent drought (D2) is the most important factor affecting metabolomic profiles and morpho-physiology

Our results strongly support Hypothesis 1, demonstrating that recurrent drought (D2) is the most significant factor influencing both the metabolomic composition and morphophysiological traits of *F. rubra*. This finding aligns with previous studies, which have shown that plants facing drought quickly engage in systemic metabolic and structural adjustments to maintain cellular integrity and ensure survival (A. Sharma et al., 2025). However, our study expands this understanding to the entire metabolome, revealing that cellular processes are significantly impacted by recurrent drought.

Recurrent drought (D2) induced a notable narrowing of the metabolic profile in *F. rubra,* characterized by reduced metabolite richness in both ionization modes and decreased Shannon diversity. This pattern suggests a resource-conservation strategy, where the plant selectively maintains key drought-response metabolites such as lipids, amino acids, phenolics, lactones, and terpenoids, which are vital for membrane stability, redox homeostasis, osmotic balance, and stress signaling under water-limited conditions (Itam et al., 2020; Michaletti et al., 2018). However, this metabolic narrowing may come at the expense of growth and defense-related metabolites, potentially lowering the plant’s long-term resilience and capacity to recover following repeated drought cycles.

Morphological and physiological traits exhibited treatment-specific responses, with recurrent drought (D2) notably increasing specific leaf area (SLA), a structural adjustment that enhances light capture while minimizing carbon investment. Nevertheless, this comes with trade-offs: thinner leaves generally retain less water and are structurally weaker, which can potentially impair photosynthetic efficiency and drought tolerance (Gonzalez-Paleo & Ravetta, 2018; Sun et al., 2025; Zhou et al., 2020). This morphological economization mirrors the metabolic streamlining observed under D2, where reduced richness and diversity reflect a focus on essential survival pathways. Together, these responses suggest that F. rubra adapts an efficient resource-use strategy under recurrent drought, but one that might compromise resilience during extended or unpredictable stress.

### 4.2 Transgenerational MeJA and Recurrent Drought Memory Reprogram Plant Metabolome and Morpho-Physiology

In line with Hypothesis 2, our findings show that drought (D1) and MeJA (M1) memories independently shaped metabolomic and morpho-physiological traits in *Festuca rubra*. D1 reduced diversity and imposed structural trade-offs, whereas M1 significantly increased Shannon diversity and Hill evenness but had no effect on richness, while also stabilizing water status and photosynthetic performance.

MeJA memory caused a distinct shift in the plant’s metabolome, shown by higher Shannon diversity and Hill Evenness (Figure S2, Table S2). This indicates a more consistent and functionally adaptable biochemical profile without increasing overall richness. The shift in metabolite levels suggests that M1 improves biochemical resilience by adjusting existing pathways, allowing for more efficient and targeted responses to recurring drought. Physiological changes in M1-treated plants closely matched these metabolomic patterns: lower specific leaf area (SLA) indicates investment in robust tissues, while higher relative water content (RWC), chlorophyll levels, and fluorescence suggest better hydration and photosynthetic performance. These trait patterns align with known effects of MeJA and other priming hormones (Bhatt et al., n.d.-b; Matkowski & Daszkowska–Golec, 2025), but our study is the first to show their lasting impact in clonal offspring never directly exposed to MeJA. This points to a transgenerational physiological imprint of hormonal priming. Interestingly, the stability in physiological traits combined with increased Shannon diversity and evenness highlights a coordinated, transgenerational reprogramming approach, one that relies on redistribution and balance, not on expanding metabolism, to improve drought resilience. While richness often dominates metabolomics research, recent studies demonstrate that Shannon diversity and evenness better reflect ecological function and stress adaptability (Petrén et al., 2024; Salgado et al., 2023). Our findings emphasize the often-overlooked significance of these distribution-focused metrics in stress memory and introduce metabolomic evenness as a new, heritable trait influenced by phytohormonal legacy.

Drought memory (D1) led to a clear reduction in Shannon diversity, without changes in richness or uniqueness, indicating a less balanced distribution of metabolites across the biochemical network. This narrowing suggests that prior drought exposure constrained the metabolic response in clonal offspring, concentrating activity into fewer dominant pathways. During recurrent drought, such an imbalance likely compromises the plant’s ability to flexibly adjust to stress, increasing reliance on a limited set of metabolic strategies. This understanding is supported by the physiological shifts observed in D1-treated plants: increased specific leaf area (SLA), reductions in relative water content (RWC), and chlorophyll levels all indicative of thinner tissues, weaker water retention, and lower photosynthetic potential. These traits are well-established indicators of drought sensitivity and resource-acquisitive strategies under stress (Blumenthal et al., 2020; Maxwell & Johnson, 2000; Perez-Harguindeguy et al., 2013b). While much of the literature frames transgenerational drought stress memory as beneficial (Avramova, 2019b; Crisp et al., 2016a), our results point to a more nuanced outcome: some stress legacies may constrain rather than enhance resilience. Specifically, we highlight a largely overlooked dimension of recurrent drought memory, that the transgenerational reduction in metabolomic diversity, accompanied by corresponding structural vulnerabilities under stress. This expands the current understanding of stress memory beyond adaptive priming, suggesting that inherited metabolic imbalance may also underlie reduced performance in clonal systems

### 4.3 Interactive Effects of MeJA Memory, Drought Memory, and Current Drought Shape Plant Responses

In line with Hypothesis 3, our findings reveal that transgenerational memories drought-induced (D1) and MeJA-induced (M1) interact in complex, context-dependent ways with recurring drought (D2) to shape both metabolomic and morpho-physiological responses in *F.rubra*.

The interaction between drought memory (D1) and recurrent drought (D2) significantly reduced metabolite richness, suggesting a form of metabolic streamlining where prior exposure to drought alters how future stress is managed biochemically. Rather than diversifying metabolic outputs in anticipation of stress, plants with drought memory appear to consolidate metabolic activity into a more restricted set of pathways under recurrent drought. This aligns with findings that repeated drought can induce energy-conserving adjustments in metabolite allocation (Ashrafi et al., 2018; Dias et al., 2021; Fàbregas & Fernie, 2019; Mukarram et al., 2021)) and that prior stress can reprogram metabolic setpoints (Avramova, 2019b). However, while metabolic efficiency is often beneficial, our results suggest that D1 may amplify the narrowing effect of D2, leading to reduced biochemical flexibility. This form of memory × stress interaction remains rarely characterized at the metabolomic level, particularly in clonal plants. By demonstrating that D1 enhances the streamlining effect of D2, we highlight a novel route by which transgenerational memory can shape, and potentially limit, metabolic responsiveness to recurring drought.

Interaction between drought memory (D1) and MeJA memory (M1) revealed that M1 effectively restored Shannon diversity and metabolomic composition in D1-treated plants, acting as a compensatory mechanism that counterbalances the metabolic narrowing imposed by drought memory. This recovery suggests that MeJA memory not only primes stress responses but also reactivates or stabilizes biosynthetic pathways suppressed by prior recurrent drought, facilitating more coordinated metabolic regulation. Although MeJA priming is well-known for enhancing abiotic stress tolerance (Avramova, 2019b; Lee et al., 2024; Zeng et al., 2024), its capacity to modulate existing stress memory effects, essentially reshaping the legacy of previous stress, is much less explored in plant biology. Recent advances in epigenetic and hormonal memory studies suggest potential cross-talk among stress memories (Chang et al., 2020; Kinoshita & Seki, 2014; Mozgova et al., 2019), but observed evidence for such interactions at the metabolome level remains scarce. By illustrating an interaction in which MeJA memory mitigates drought memory’s restrictive impacts, our findings introduce a novel dimension of memory interplay, where one form of stress legacy actively influences another, highlighting a previously underappreciated layer of metabolic plasticity and resilience.

The interaction between MeJA memory (M1) and recurrent drought (D2) further underscores M1’s role in maintaining metabolic coherence. While D2 alone increased metabolomic uniqueness reflecting greater variability and divergence in biochemical regulation this effect was notably tempered when M1 was present. Uniqueness was lowest in M1-only plants, and its addition under D2 conditions reduced the otherwise heightened differentiation in metabolic profiles. This suggests that MeJA memory may constrain overactive or disorganized stress responses, helping to stabilize metabolic networks under fluctuating environmental conditions. Although MeJA is well-recognized for enhancing preparedness to abiotic stress (Hewedy et al., 2023b; Wasternack & Strnad, 2019; Yu et al., 2019), its role in buffering metabolic divergence during acute drought has received limited attention. Our findings highlight that M1 not only primes stress pathways but also modulates the amplitude and variability of drought-induced metabolic shifts, supporting a role for MeJA memory as a metabolic stabilizer that reinforces coordination and predictability in plant responses to environmental stress.

Another significant finding was the emergence of a strong three-way interaction (D1 × D2 × M1) affecting metabolomic uniqueness. Rather than producing additive effects, these memory types integrated to generate distinct, highly variable biochemical phenotypes. This reflects a previously uncharacterized layer of metabolic plasticity driven by cumulative and interacting stress legacies. To our knowledge, this is the first observed evidence demonstrating that multiple stress memories can combine to produce emergent metabolomic profiles.

At the cellular level, such memory integration likely reflects a dynamic recalibration of metabolic networks, allowing plants to integrate both historical and immediate environmental cues. Rather than returning to fixed memory states, plants appear to flexibly reconfigure their metabolomes in response to the specific sequence and combination of prior stress exposures. Adaptive plasticity is supported by emerging evidence on multi-stress memory interactions and metabolic resetting(e.g., Schwachtje et al., 2019; Szepesi & Szőllősi, 2018; Zhang et al., 2022). This flexibility may enhance cellular homeostasis and resource-use efficiency, especially under fluctuating conditions. However, in clonal populations, such reprogramming could also lead to increased phenotypic divergence among genetically identical individuals, a phenomenon previously linked to epigenetic variation and stress-induced metabolic noise (Crisp et al., 2016b). Our findings suggest that memory crosstalk at the metabolic level introduces both stabilizing and diversifying forces, which together shape how plants navigate environmental unpredictability over time.

Morpho-physiological data further support the stabilizing role of MeJA memory (M1). Plants with M1 treatments consistently maintained higher relative water content and chlorophyll levels, without exhibiting structural trade-offs such as increased specific leaf area (SLA)—a pattern suggesting more integrated and energy-efficient stress responses. In contrast, the D1 × D2 combination was associated with reduced coordination between metabolic and morphological traits, indicating a fragmentation of stress responses under compounded drought memory. This divergence highlights M1’s unique ability to maintain systemic coherence, even in the presence of overlapping stress legacies. Such trait coordination has been identified as a key determinant of drought resilience (Kambona et al., 2023b; Perez-Harguindeguy et al., 2013b), but its modulation by transgenerational memory remains poorly characterized. Our findings highlight the importance of studying memory crosstalk in clonal and perennial species, where sequential environmental signals can interact across generations, potentially amplifying or mitigating adaptive trade-offs.

### 4.4 Metabolome and physiology relate strongly to each other

Our findings support a tightly integrated relationship between metabolomic and morpho-physiological traits, reinforcing Hypothesis 4 (H4). Traits such as specific leaf area (SLA), relative water content (RWC), and chlorophyll content (CCS2) emerged as strong predictors of metabolomic variation, reflecting coordinated biochemical and structural adaptation to both current drought and stress legacies (e.g., Barrs & Weatherley, 1962; Künzi et al., 2025; Li et al., 2024; Maxwell & Johnson, 2000; Pérez-Ramos et al., 2013) .

Crucially, MeJA memory (M1) acted as a stabilizing force across both domains. M1-treated plants maintained higher RWC and chlorophyll levels while avoiding elevated SLA, indicative of more efficient resource use and structural conservation. These physiological advantages aligned with targeted upregulation of protective metabolites, particularly flavonoids, curcuminoids, lipids, and terpenoids known to enhance osmotic regulation, antioxidative defense, and membrane integrity (Fàbregas & Fernie, 2019; Falcone Ferreyra et al., 2012).These metabolites were enriched in M1D2 plants, reflecting memory-mediated biochemical readiness under stress.

In contrast, plants experiencing only drought memories (D1, D2) or their combination (D1D2) exhibited elevated SLA and accumulation of energetically costly metabolites like phosphocholines and ceramides, suggesting less targeted responses and reduced physiological coordination. Notably, M1 suppressed these broad-spectrum responses when co-applied with D2, indicating a strategy of metabolic economy and functional specificity (Crisp et al., 2016b; M. Sharma et al., 2022).

Altogether, these findings demonstrate that MeJA memory not only modulates stress responses at the metabolomic level but also aligns them with morpho-physiological performance. This dual-level optimization highlights a core mechanism by which priming enhances drought resilience: by coordinating structural traits with efficient, targeted metabolic programming. Such integration is especially relevant in clonal systems, where compounded stress legacies require precise, energy-efficient adaptation strategies.

## 5. Conclusion

This study presents the first comprehensive evidence that multiple transgenerational stress memories, specifically drought and methyl jasmonate (MeJA) memory, interact in synergistic, non-additive ways to shape both metabolomic and morphophysiological traits in a clonal grass. We show that MeJA memory functions as a central modulator, restoring diversity, dampening metabolic divergence, and aligning physiological performance under recurrent drought. Notably, we demonstrate that MeJA can modulate the expression of other memory types, introducing a novel concept of memory crosstalk in plant stress biology.

Beyond model plant systems, our findings offer practical insights into how hormonal priming, particularly MeJA priming, can be harnessed to enhance drought tolerance across diverse clonal and perennial species. The identification of coordinated metabolic and structural responses driven by priming in clonal offspring underscores the potential of integrated priming strategies to enhance long-term resilience. Overall, our work advances our understanding of plant stress memory and opens new avenues for harnessing biochemical legacies to improve drought adaptation in both ecological and agricultural systems.

## Data Availability

The data supporting the findings of this study are currently held by the authors and will be deposited in the Zenodo repository upon acceptance of the manuscript.

## Acknowledgements

GAUK project number 171724 supported the study. It was also partly supported by GAČR 22-00761S, a long-term research development project no. RVO 67985939 of the Czech Academy of Sciences and institutional support for science and research of the Ministry of Education, Youth, and Sports of the Czech Republic.

## Author contributions

TB, DT, and ZM conceptualized the study; TB, ZM, and DT did the experimental setup; NR and TB meticulously prepared the samples for UPLC analysis, while NR, DT, and JS optimized the UPLC-MS method; JS and TC performed UPLC-MS analysis; The raw data were processed to obtain metabolite feature data by TB, DT, and JS; TB carried out data analysis with the help of DT and ZM. TB led the writing, with in-depth editing and suggestions from DT and ZM; DT and ZM are the senior authors. All authors approved the submission and declare no conflict of interest.

## Supplementary

**Table S1:**
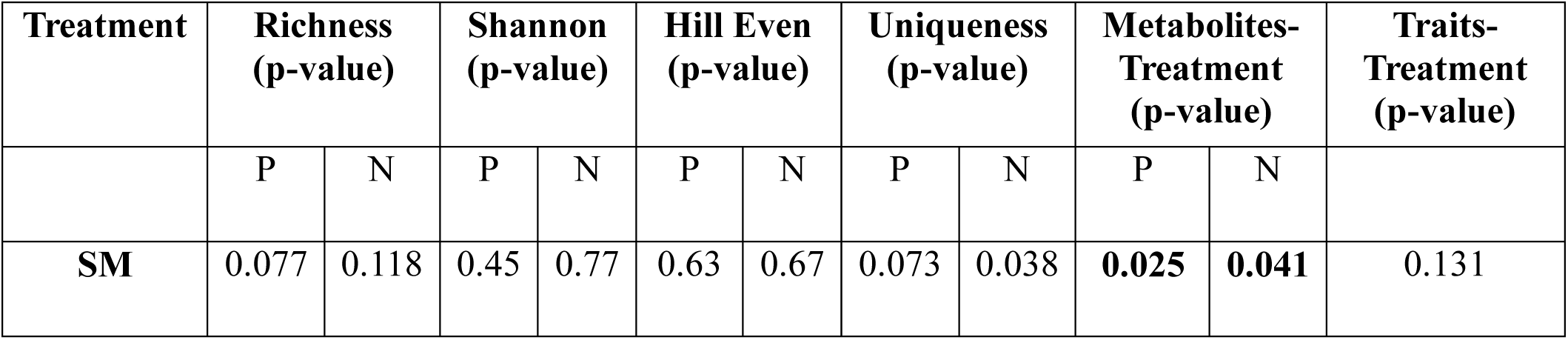
This table presents *p*-values comparing metabolome diversity metrics and metabolomic/trait-level responses for soil memory (SM). Metabolome analysis was performed in both positive (P) and negative (N) ionization modes. Significant results (*p* < 0.05) indicate treatment-induced effects and are in bold.

**Text S2: Experimental design**

The experiment represented a factorial cross of drought and MeJA treatments, each with two levels (yes/no). Each treatment had 10 replicates. The soil utilized in the experiment was sourced from the same meadow where the experimental plants were originally collected. Pots with a volume of 500 ml were filled with the soil, and one *F. rubra* ramet was planted into each pot. All the planted ramets were of similar age and size (approximately 3 weeks old and 3.5 to 4 cm tall). The plants were placed in a growth chamber with long-day conditions, where the temperature ranged from 17 to 20 °C during the day (16 hours) and 7 to 10 °C at night (8 hours). These conditions reflect conditions at the original locality during peak growing season. The plants were allowed to establish for 21 days before starting the experiment. The experiment included cycles of drought and recovery phases, each lasting 30 days (Drought stage 1 - Recovery stage 1 - Drought stage 2 - Recovery stage 2 - Drought stage 3 - Recovery stage 3).

Control watering treatment received regular watering daily, maintaining 1 cm of water above the pot base in the trays throughout the experiment. Drought-stressed plants received 50 ml of water in specific watering intervals during the drought stages: every 7th day during drought stage 1, every 10th day in drought stage 2, and every 15th day in drought stage 3. There was a progressive increase in drought stress among phases because treating plants to an excessive amount of stress initially can result in total plant loss, forcing the experiment to terminate after the first drought stage. In each phase, we attempted to expose the plants to as intense drought stress as possible but made sure they would survive successive drought stages. During the recovery stage, drought-stressed plants received the same watering as the control ones. The soil moisture has been monitored using TMS dataloggers (Wild et al., 2019).MeJA-treated *F. rubra* plants received 5 ml of 10 µM MeJA by pouring it directly onto the soil every 24 hours during the stress phases, but not during recovery. Only drought plants also got 5 ml of distilled water when 5 ml of 10 µM MeJA was given to MeJA-treated plants during drought. Previous studies predominantly relied on foliar application of MeJA for plant treatment (e.g. Tayyab et al., 2020; Chungloo et al., 2023)We decided to apply it to the soil as foliar application has various disadvantages. The volatile nature of MeJA poses a risk of environmental loss when applied through foliar application, and there is an additional concern of uneven MeJA distribution among leaves across the plant. Therefore, soil application is a viable solution to address these challenges effectively. Soil is an efficient absorber, thus minimizing the risk of MeJA loss to the environment, while root uptake guarantees a more consistent distribution of MeJA throughout the entire plant.

**Figure S1:**
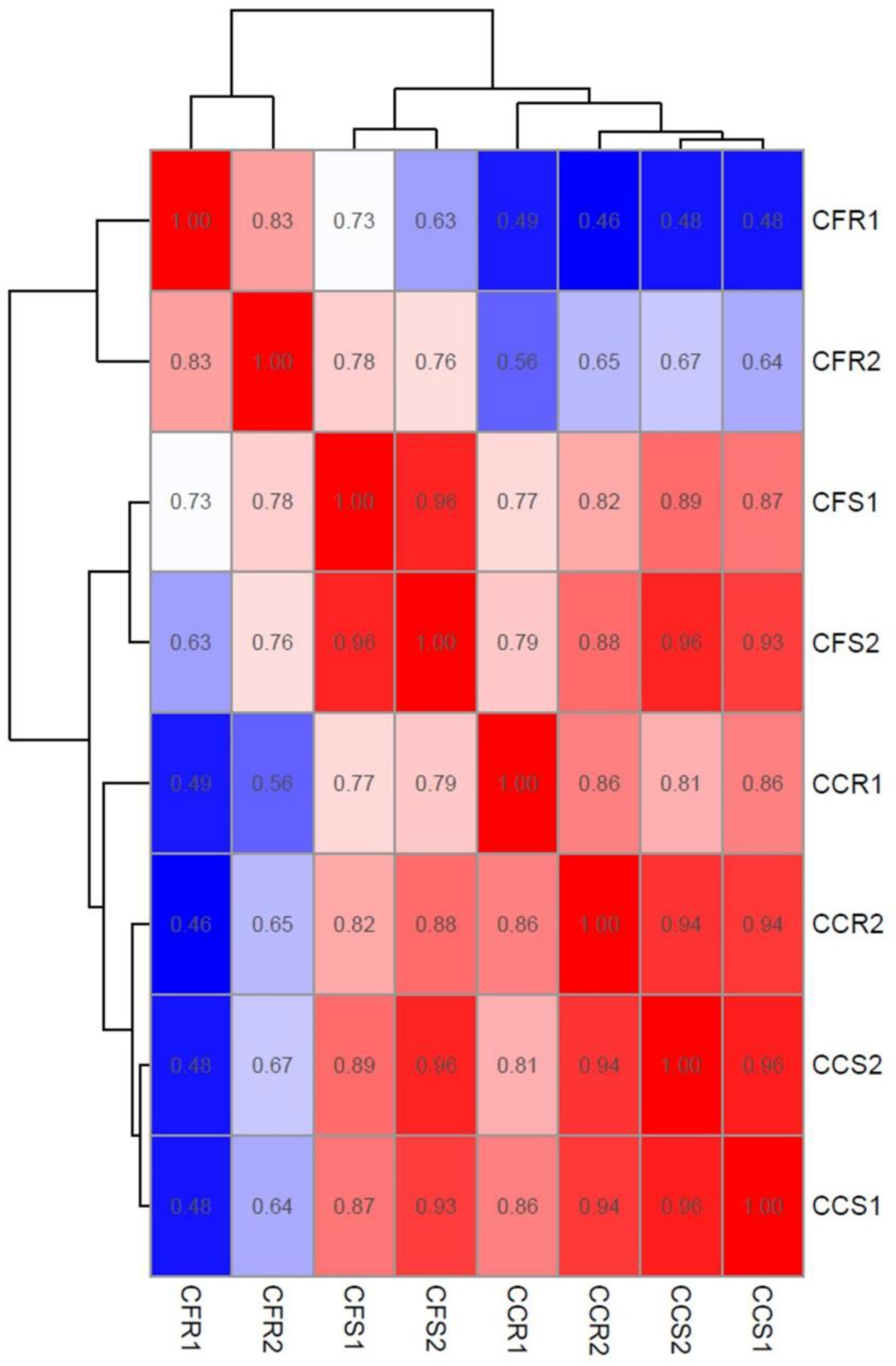
Correlation heatmap showing the relationship among chlorophyll fluorescence parameters measured during the stress phase (CFS1, CFS2), recovery phase (CFR1, CFR2), chlorophyll content measured during the stress phase (CCS1, CCS2), and recovery phase (CCR1, CCR2). Correlation coefficients are color-coded from dark blue (low correlation) to dark red (high correlation), with values displayed within each cell.

**Table S2:** Table showing effects of M1, D1, D2, i.e., MeJA memory, drought memory, and current drought respectively, and their interactions on Hill Eveness. The variation explained is shown for the significant predictors (p < 0.05).

| Treatment | Hill Eveness |  |  |  |
| --- | --- | --- | --- | --- |
|  | Positive mode |  | Negative mode |  |
|  | P-value | Variation explained | P-value | Variation explained |
| D1 | 0.01 | 10.66 | 0.97 | - |
| D2 | 0.357 | - | 0.01 | 14.35 |
| M1 | 0.007 | 19.78 | 0.37 | - |
| D1×D2 | 0.8 | - | 0.43 | - |
| D1×M1 | 0.011 | 10.47 | 0.36 | - |
| D2×M1 | 0.489 | - | 0.89 | - |
| D1×D2×M1 | 0.167 | - | 0.897 | - |

**Figure S2:**
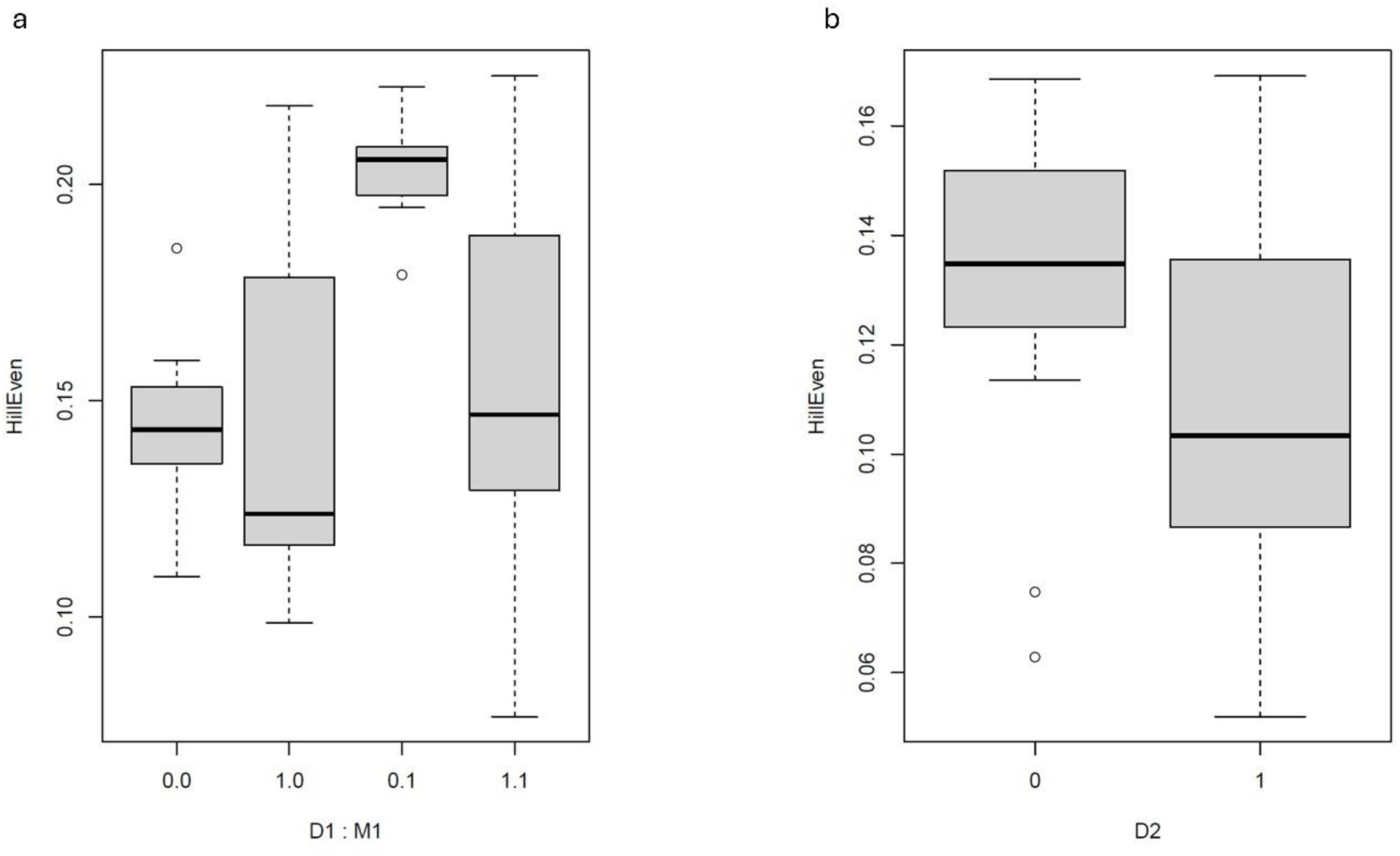
Boxplots showing the effects of interactions of M1, D1, D2, i.e. MeJA memory, drought memory and current drought, respectively in HillEven for (a) Positive mode and (b) negative mode in *F. rubra*. The X axis shows the levels of the interacting treatment, with 0 indicating absent and 1 present treatment (drought or MeJa).

**Figure S3:**
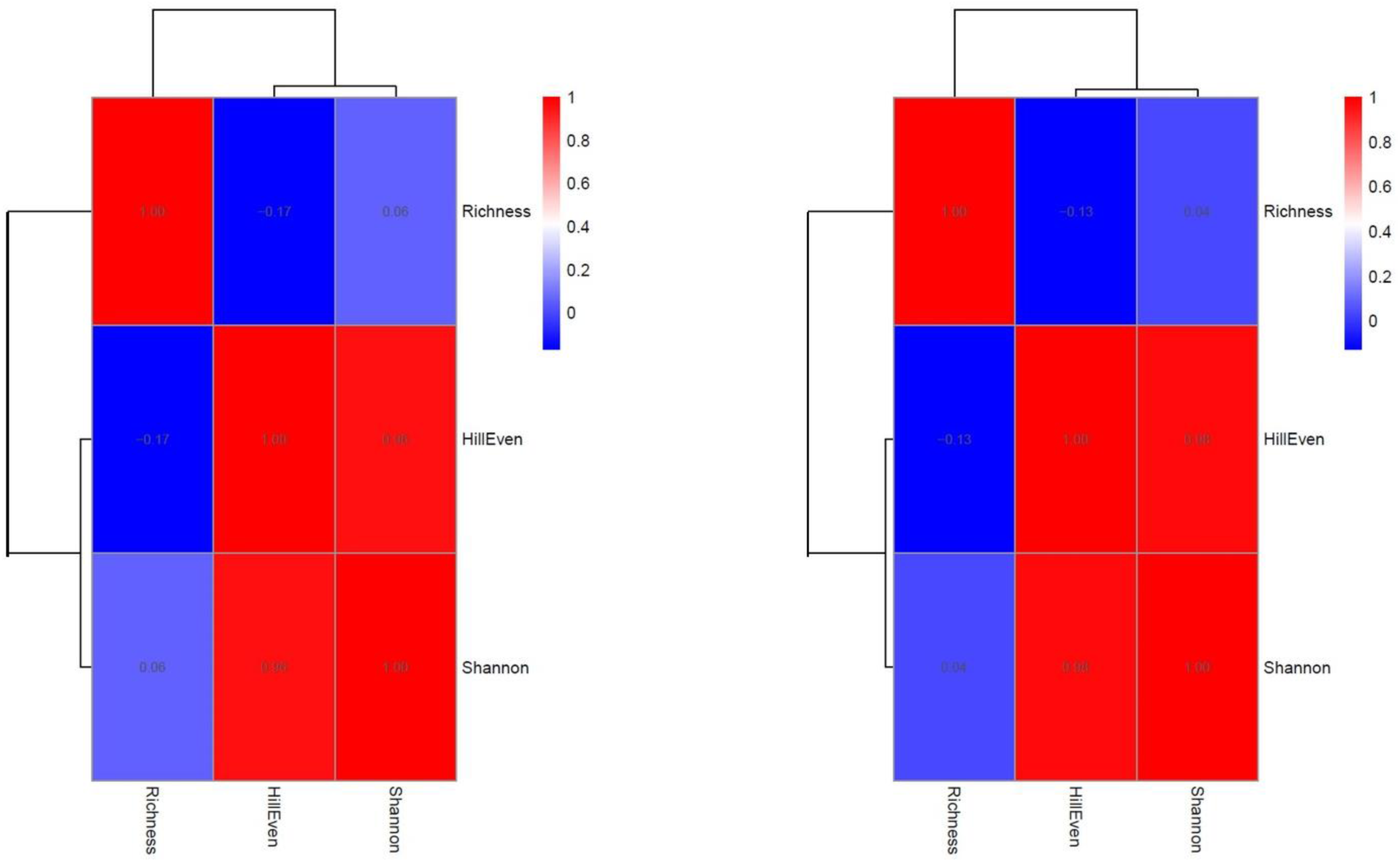
Correlation heatmap showing the relationship among Richness, Shannon diversity, and HillEvenness. Correlation coefficients are color-coded from dark blue (low correlation) to dark red (high correlation), with values displayed within each cell.

**Figure S4:**
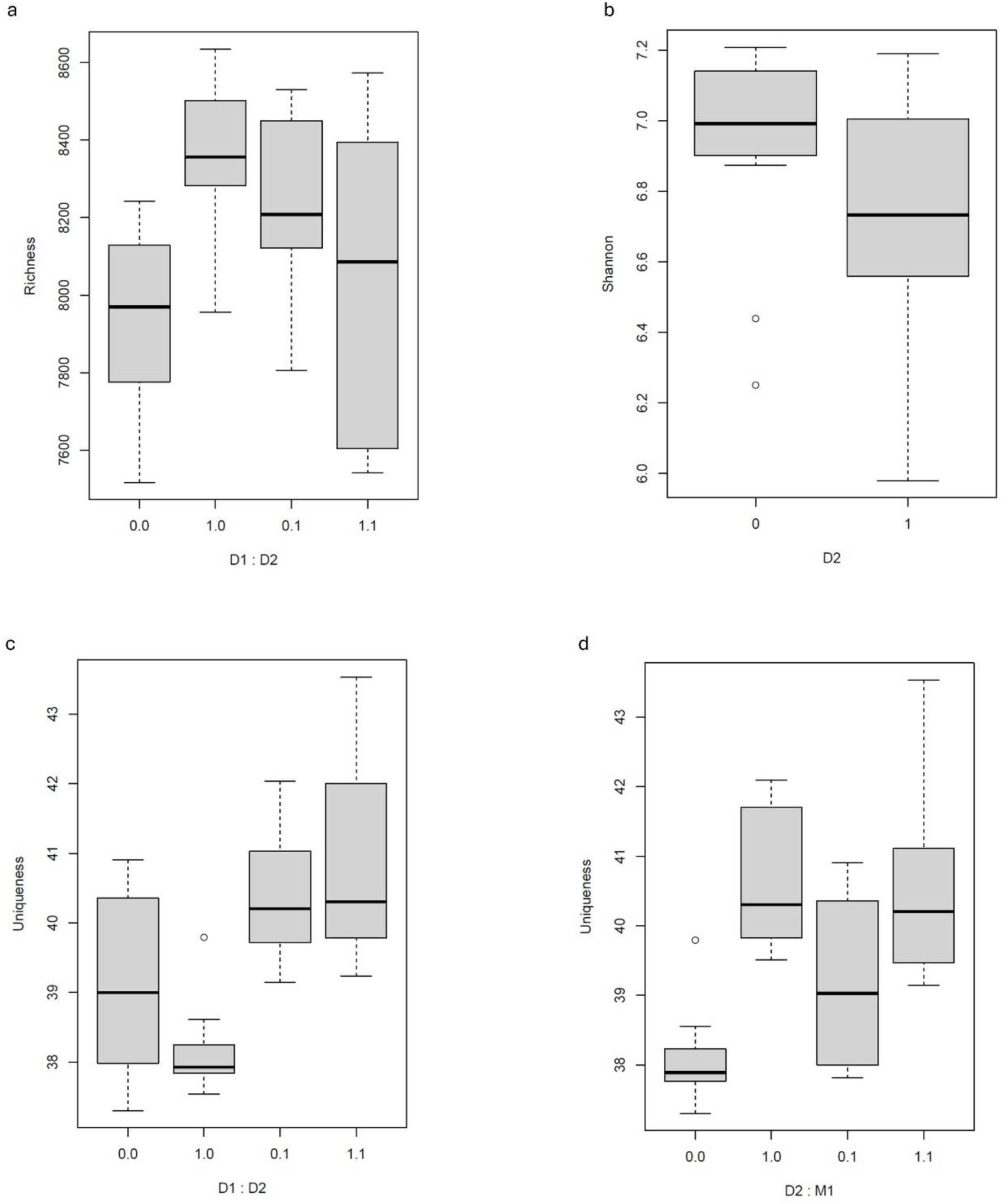
Boxplots showing the effects of interactions M1, D1, D2, i.e. MeJA memory, drought memory and current drought, respectively, in positive mode for (a) Richness, (b) Shannon diversity, and (c and d) Uniqueness in *F. rubra*. The X axis shows the levels of the interacting treatment, with 0 indicating absent and 1 present treatment (drought or MeJA).

**Figure S5:**
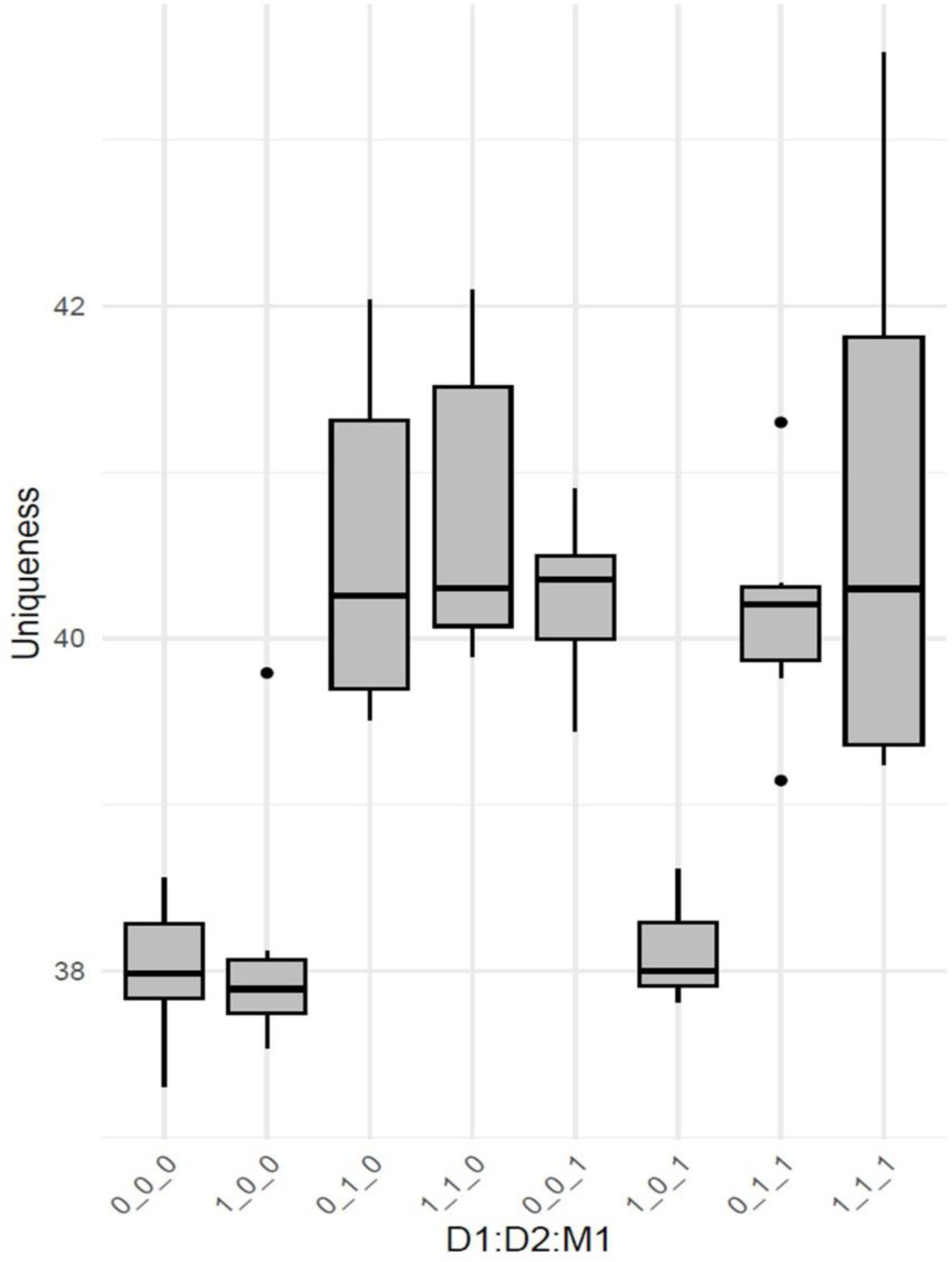
Boxplots show the distribution of metabolic uniqueness scores for each treatment group under negative ionization mode, representing the triple interaction of drought memory, current drought, and MeJA memory. In each code, a 1 in the first position indicates the presence of drought 1 (D1), a 1 in the second position indicates the presence of drought 2 (D2), and a 1 in the third position indicates the presence of the morphological modifier (M1). For example, 1_0_0 represents D1 only, 0_1_0 represents D2 only, and 1_1_1 represents a combination of D1, D2, and M1. The control group is 0_0_0, where no drought or modifier is applied.

**Figure S6:**
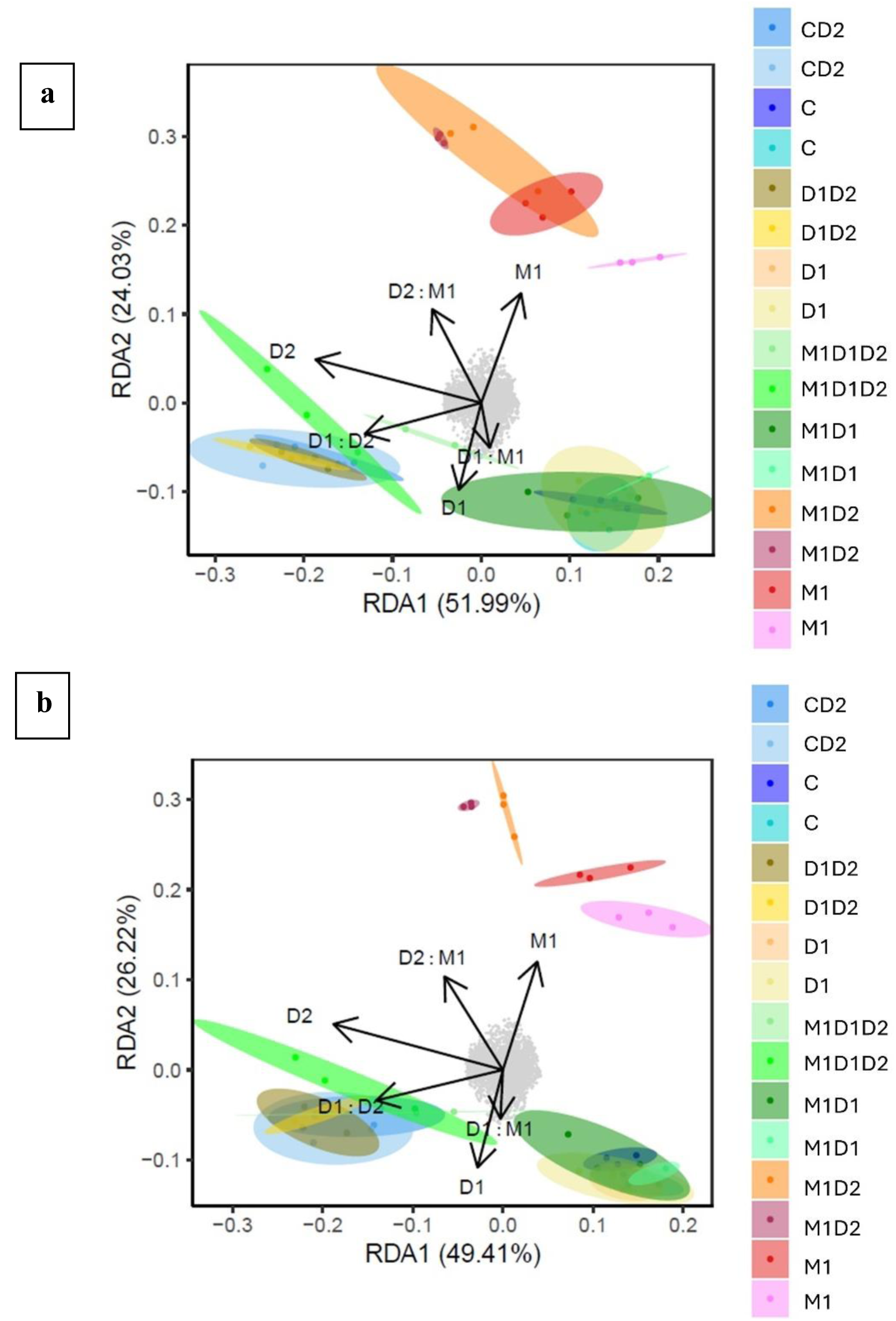
The RDA plots both ionization modes (**a**)positive and (**b**) negative, providing a clear view of the variation in metabolite profiles explained by M1, D1, D2, i.e., MeJA memory, drought memory and current drought, respectively.

**Figure S7:**
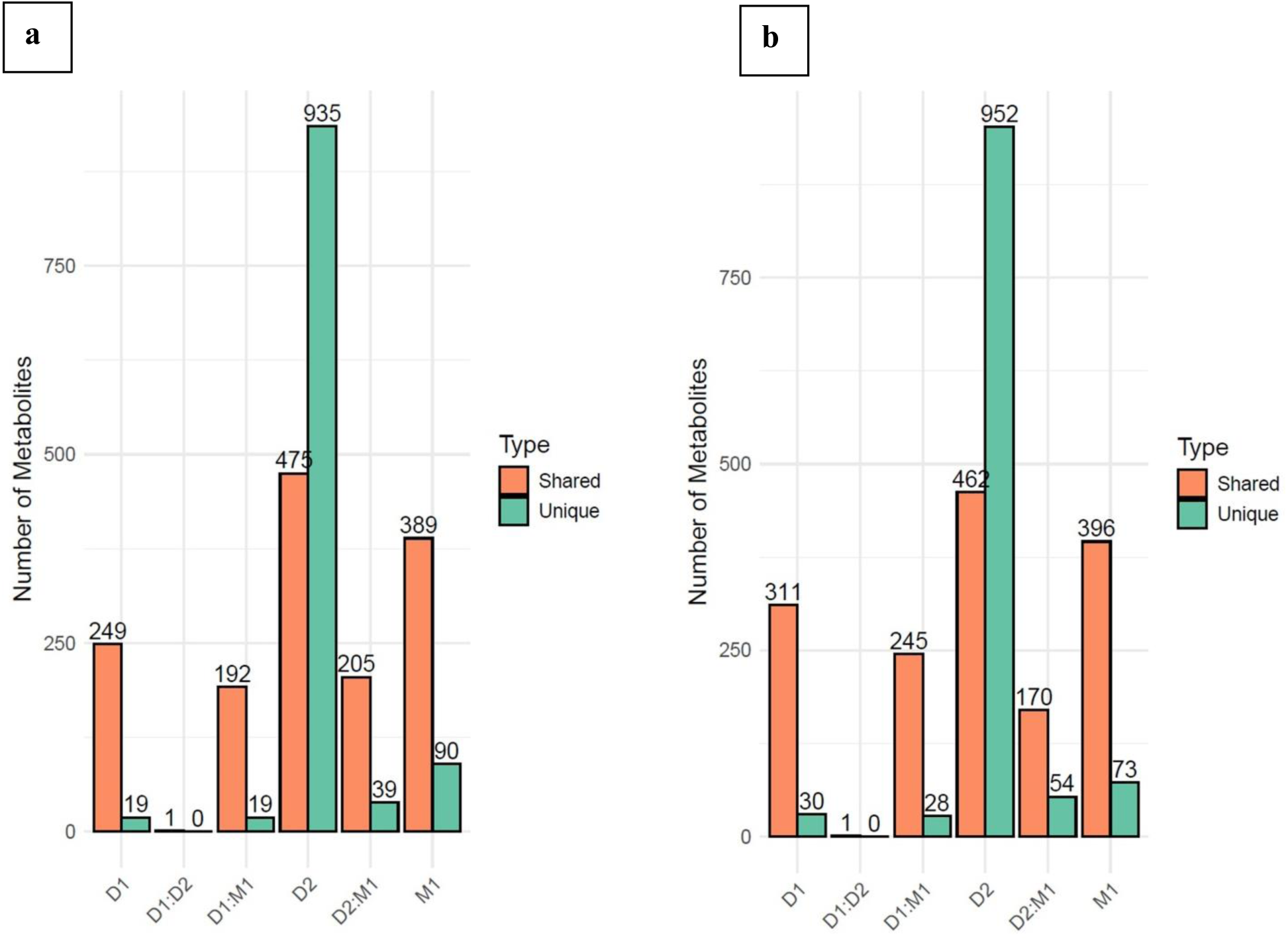
Bar plots representing the number of significant metabolites associated with drought memory (D1), MeJA memory (M1), current drought (D2), and their interactions (D1×D2, D1×M1, D2×M1) under positive ionization mode (a) and negative ionization mode (b). Each treatment or interaction is shown with two bars: shared metabolites (orange) indicate those common with other conditions, while unique metabolites (green) represent those specific to a given condition or interaction. The values above each bar indicate the exact number of metabolites in each category.

**Figure S8:**
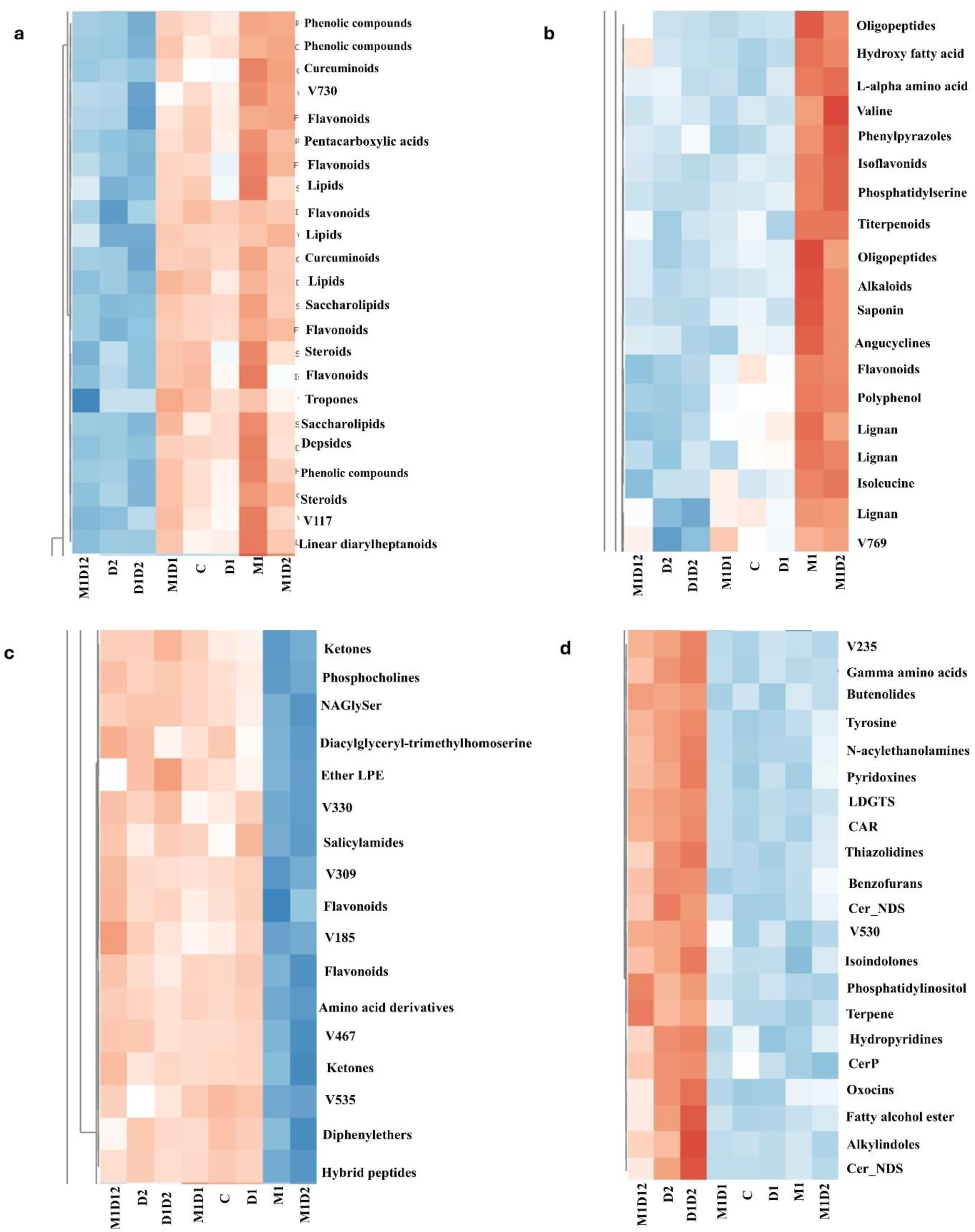
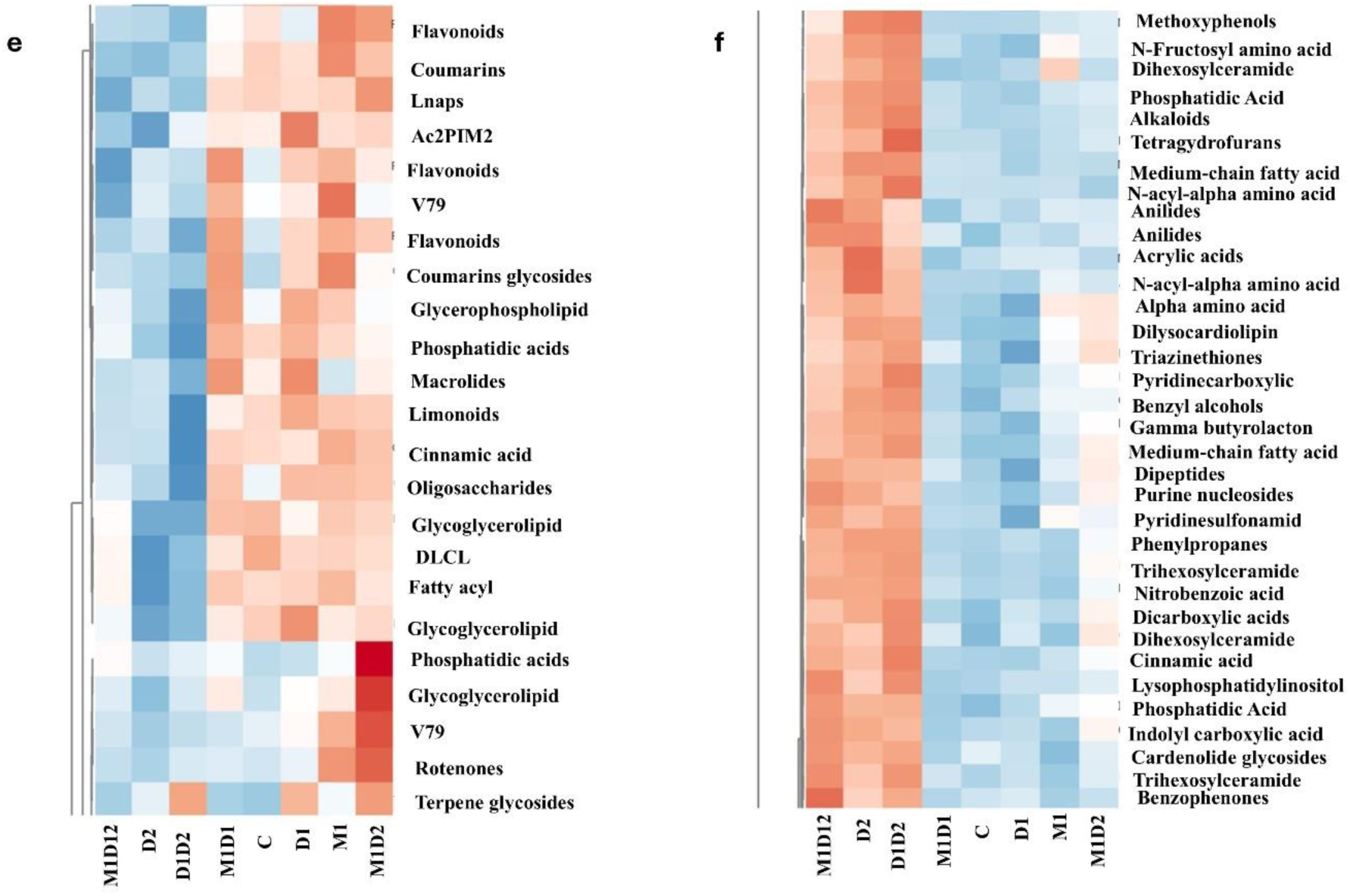
Heatmap displaying the scaled intensity values of differentially accumulated metabolites across various treatment groups in *F. rubra* (M1, D1, D2, i.e. MeJA memory, drought memory and current drought respectively), others are the plants which got combination M1, D2 and D2 treatment, while missing code in combination indicates that plants where not treated by the given treatment. C indicates Control, i.e., no drought or MeJa treatment memory, and in well-watered conditions. Heatmap a to d shows cluster 1 to 4, respectively for positive mode, while heatmap e and f shows cluster 1 and 2, respectively for negative mode

## References

Ashrafi, M., Azimi-Moqadam, M.-R., Moradi, P., MohseniFard, E., Shekari, F., & Kompany-Zareh, M. (2018). Effect of drought stress on metabolite adjustments in drought tolerant and sensitive thyme. Plant Physiology and Biochemistry, 132, 391–399. 10.1016/j.plaphy.2018.09.009

Avramova, Z. (2019a). Defence-related priming and responses to recurring drought: Two manifestations of plant transcriptional memory mediated by the ABA and JA signalling pathways. *Plant*, Cell & Environment, 42(3), 983–997.

Avramova, Z. (2019b). Defence-related priming and responses to recurring drought: Two manifestations of plant transcriptional memory mediated by the ABA and JA signalling pathways. *Plant*, Cell & Environment, 42(3), 983–997.

Barrs, H. D., & Weatherley, P. E. (1962). A re-examination of the relative turgidity technique for estimating water deficits in leaves. Australian Journal of Biological Sciences, 15(3), 413–428.

Bhatt, T., Thakur, D., & Munzbergova, Z. (n.d.-a). Unveiling the Interactive Dynamics: Recurring Drought, Methyl Jasmonate, and Plant Performance in a Perennial Grass. Methyl Jasmonate, and Plant Performance in a Perennial Grass.

Bhatt, T., Thakur, D., & Munzbergova, Z. (n.d.-b). Unveiling the Interactive Dynamics: Recurring Drought, Methyl Jasmonate, and Plant Performance in a Perennial Grass. Methyl Jasmonate, and Plant Performance in a Perennial Grass.

Björkman, O., & Demmig, B. (1987). Photon yield of O 2 evolution and chlorophyll fluorescence characteristics at 77 K among vascular plants of diverse origins. Planta, 170, 489–504.

Blumenthal, D. M., Mueller, K. E., Kray, J. A., Ocheltree, T. W., Augustine, D. J., & Wilcox, K. R. (2020). Traits link drought resistance with herbivore defence and plant economics in semi-arid grasslands: The central roles of phenology and leaf dry matter content. Journal of Ecology, 108(6), 2336–2351.

Calderón-Santiago, M., Fernández-Peralbo, M. A., Priego-Capote, F., & Luque de Castro, M. D. (2016). MSCombine: a tool for merging untargeted metabolomic data from high-resolution mass spectrometry in the positive and negative ionization modes. Metabolomics, 12(3), 43. 10.1007/s11306-016-0970-4

Chang, Y.-N., Zhu, C., Jiang, J., Zhang, H., Zhu, J.-K., & Duan, C.-G. (2020). Epigenetic regulation in plant abiotic stress responses. Journal of Integrative Plant Biology, 62(5), 563–580. 10.1111/jipb.12901

Chungloo, D., Tisarum, R., Sotesaritkul, T., Praseartkul, P., Himanshu, S. K., Datta, A., & Cha-um, S. (2023). Exogenous Foliar Application of Methyl Jasmonate Alleviates Water-Deficit Stress in Andrographis paniculata. Journal of Soil Science and Plant Nutrition. 10.1007/s42729-023-01414-0

Crisp, P. A., Ganguly, D., Eichten, S. R., Borevitz, J. O., & Pogson, B. J. (2016a). Reconsidering plant memory: Intersections between stress recovery, RNA turnover, and epigenetics. Science Advances, 2(2), e1501340.

Crisp, P. A., Ganguly, D., Eichten, S. R., Borevitz, J. O., & Pogson, B. J. (2016b). Reconsidering plant memory: Intersections between stress recovery, RNA turnover, and epigenetics. Science Advances, 2(2), e1501340.

Dias, M. C., Pinto, D. C. G. A., Figueiredo, C., Santos, C., & Silva, A. M. S. (2021). Phenolic and lipophilic metabolite adjustments in Olea europaea (olive) trees during drought stress and recovery. Phytochemistry, 185, 112695. 10.1016/j.phytochem.2021.112695

Fàbregas, N., & Fernie, A. R. (2019). The metabolic response to drought. Journal of Experimental Botany, 70(4), 1077–1085. 10.1093/jxb/ery437

Falcone Ferreyra, M. L., Rius, S. P., & Casati, P. (2012). Flavonoids: biosynthesis, biological functions, and biotechnological applications. Frontiers in Plant Science, 3, 222.

Gonzalez-Paleo, L., & Ravetta, D. A. (2018). Relationship between photosynthetic rate, water use and leaf structure in desert annual and perennial forbs differing in their growth. Photosynthetica, 56(4), 1177–1187. 10.1007/s11099-018-0810-z

Hewedy, O. A., Elsheery, N. I., Karkour, A. M., Elhamouly, N., Arafa, R. A., Mahmoud, G. A.-E., Dawood, M. F.-A., Hussein, W. E., Mansour, A., & Amin, D. H. (2023a). Jasmonic acid regulates plant development and orchestrates stress response during tough times. Environmental and Experimental Botany, 208, 105260.

Hewedy, O. A., Elsheery, N. I., Karkour, A. M., Elhamouly, N., Arafa, R. A., Mahmoud, G. A.-E., Dawood, M. F.-A., Hussein, W. E., Mansour, A., & Amin, D. H. (2023b). Jasmonic acid regulates plant development and orchestrates stress response during tough times. Environmental and Experimental Botany, 208, 105260.

Hoover, D. L., Pfennigwerth, A. A., & Duniway, M. C. (2021). Drought resistance and resilience: The role of soil moisture–plant interactions and legacies in a dryland ecosystem. Journal of Ecology, 109(9), 3280–3294.

Itam, M., Mega, R., Tadano, S., Abdelrahman, M., Matsunaga, S., Yamasaki, Y., Akashi, K., & Tsujimoto, H. (2020). Metabolic and physiological responses to progressive drought stress in bread wheat. Scientific Reports, 10(1), 17189. 10.1038/s41598-020-74303-6

Jiao, T., Williams, C. A., De Kauwe, M. G., & Medlyn, B. E. (2023a). Limited Evidence of Cumulative Effects From Recurrent Droughts in Vegetation Responses to Australia’s Millennium Drought. Journal of Geophysical Research: Biogeosciences, 128(5), e2022JG006818. 10.1029/2022JG006818

Jiao, T., Williams, C. A., De Kauwe, M. G., & Medlyn, B. E. (2023b). Limited evidence of cumulative effects from recurrent droughts in vegetation responses to Australia’s Millennium Drought. Journal of Geophysical Research: Biogeosciences, 128(5), e2022JG006818.

Kambona, C. M., Koua, P. A., Léon, J., & Ballvora, A. (2023a). Stress memory and its regulation in plants experiencing recurrent drought conditions. Theoretical and Applied Genetics, 136(2), 26.

Kambona, C. M., Koua, P. A., Léon, J., & Ballvora, A. (2023b). Stress memory and its regulation in plants experiencing recurrent drought conditions. Theoretical and Applied Genetics, 136(2), 26. 10.1007/s00122-023-04313-1

Kinoshita, T., & Seki, M. (2014). Epigenetic Memory for Stress Response and Adaptation in Plants. Plant and Cell Physiology, 55(11), 1859–1863. 10.1093/pcp/pcu125

Künzi, Y., Zeiter, M., Fischer, M., & Stampfli, A. (2025). Rooting depth and specific leaf area modify the impact of experimental drought duration on temperate grassland species. Journal of Ecology, 113(2), 445–458. 10.1111/1365-2745.14468

Lee, D. Y., Park, D. Y., & Park, H. C. (2024). Exogenous methyl jasmonate mediates tolerance of heat stress in Korean fir (Abies koreana). Plant Biotechnology Reports, 18(4), 525–534. 10.1007/s11816-024-00912-6

Li, Z.-K., Li, H.-L., Gong, X.-W., Wang, H.-F., & Hao, G.-Y. (2024). Prediction and mapping of leaf water content in Populus alba var. pyramidalis using hyperspectral imagery. Plant Methods, 20(1), 184. 10.1186/s13007-024-01312-1

Liu, L., Cao, X., Zhai, Z., Ma, S., Tian, Y., & Cheng, J. (2022). Direct evidence of drought stress memory in mulberry from a physiological perspective: Antioxidative, osmotic and phytohormonal regulations. Plant Physiology and Biochemistry, 186, 76–87.

Luo, W., Ma, W., Song, L., Te, N., Chen, J., Muraina, T. O., Wilkins, K., Griffin-Nolan, R. J., Ma, T., & Qian, J. (2023). Compensatory dynamics drive grassland recovery from drought. Journal of Ecology, 111(6), 1281–1291.

Lv, P., Sun, S., Li, Y., Zhao, S., Zhang, J., Hu, Y., Yue, P., & Zuo, X. (2024). Growing-season drought and nitrogen addition interactively impair grassland ecosystem stability by reducing species diversity, asynchrony, and stability. Science of The Total Environment, 912, 169122.

Matkowski, H., & Daszkowska–Golec, A. (2025). Wisdom comes after facts – An update on plants priming using phytohormones. Journal of Plant Physiology, 305, 154414. 10.1016/j.jplph.2024.154414

Maxwell, K., & Johnson, G. N. (2000). Chlorophyll fluorescence—a practical guide. Journal of Experimental Botany, 51(345), 659–668.

Michaletti, A., Naghavi, M. R., Toorchi, M., Zolla, L., & Rinalducci, S. (2018). Metabolomics and proteomics reveal drought-stress responses of leaf tissues from spring-wheat. Scientific Reports, 8(1), 5710.

Mohamed, H. I., & Latif, H. H. (2017). Improvement of drought tolerance of soybean plants by using methyl jasmonate. Physiology and Molecular Biology of Plants, 23(3), 545–556.

Möhl, P., Vorkauf, M., Kahmen, A., & Hiltbrunner, E. (2023). Recurrent summer drought affects biomass production and community composition independently of snowmelt manipulation in alpine grassland. Journal of Ecology, 111(11), 2357–2375.

Mozgova, I., Mikulski, P., Pecinka, A., & Farrona, S. (2019). Epigenetic Mechanisms of Abiotic Stress Response and Memory in Plants. In R. Alvarez-Venegas, C. De-la-Peña, & J. A. Casas-Mollano (Eds.), Epigenetics in Plants of Agronomic Importance: Fundamentals and Applications: Transcriptional Regulation and Chromatin Remodelling in Plants (pp. 1–64). Springer International Publishing. 10.1007/978-3-030-14760-0_1

Mukarram, M., Choudhary, S., Kurjak, D., Petek, A., & Khan, M. M. A. (2021). Drought: Sensing, signalling, effects and tolerance in higher plants. Physiologia Plantarum, 172(2), 1291–1300. 10.1111/ppl.13423

Muller, B., Pantin, F., Génard, M., Turc, O., Freixes, S., Piques, M., & Gibon, Y. (2011). Water deficits uncouple growth from photosynthesis, increase C content, and modify the relationships between C and growth in sink organs. Journal of Experimental Botany, 62(6), 1715–1729. 10.1093/jxb/erq438

Nordström, A., Want, E., Northen, T., Lehtiö, J., & Siuzdak, G. (2008). Multiple Ionization Mass Spectrometry Strategy Used To Reveal the Complexity of Metabolomics. Analytical Chemistry, 80(2), 421–429. 10.1021/ac701982e

Pang, Z., Lu, Y., Zhou, G., Hui, F., Xu, L., Viau, C., Spigelman, A. F., MacDonald, P. E., Wishart, D. S., Li, S., & Xia, J. (2024). MetaboAnalyst 6.0: towards a unified platform for metabolomics data processing, analysis and interpretation. Nucleic Acids Research, 52(W1), W398–W406. 10.1093/nar/gkae253

Perez-Harguindeguy, N., Diaz, S., Garnier, E., Lavorel, S., Poorter, H., Jaureguiberry, P., Bret-Harte, M. S., Cornwell, W. K., Craine, J. M., & Gurvich, D. E. (2013a). New handbook for standardised measurement of plant functional traits worldwide. Aust. Bot. 61, 167–234.

Perez-Harguindeguy, N., Diaz, S., Garnier, E., Lavorel, S., Poorter, H., Jaureguiberry, P., Bret-Harte, M. S., Cornwell, W. K., Craine, J. M., & Gurvich, D. E. (2013b). New handbook for standardised measurement of plant functional traits worldwide. Aust. Bot. 61, 167–234.

Pérez-Ramos, I. M., Volaire, F., Fattet, M., Blanchard, A., & Roumet, C. (2013). Tradeoffs between functional strategies for resource-use and drought-survival in Mediterranean rangeland species. Environmental and Experimental Botany, 87, 126–136.

Petrén, H., Anaia, R. A., Aragam, K. Sen, Bräutigam, A., Eckert, S., Heinen, R., Jakobs, R., Ojeda-Prieto, L., Popp, M., Sasidharan, R., Schnitzler, J.-P., Steppuhn, A., Thon, F. M., Unsicker, S. B., van Dam, N. M., Weisser, W. W., Wittmann, M. J., Yepes, S., Ziaja, D.,… Junker, R. R. (2024). Understanding the chemodiversity of plants: Quantification, variation and ecological function. Ecological Monographs, 94(4), e1635. 10.1002/ecm.1635

Petrén, H., Köllner, T. G., & Junker, R. R. (2023). Quantifying chemodiversity considering biochemical and structural properties of compounds with the R package chemodiv. New Phytologist, 237(6), 2478–2492. 10.1111/nph.18685

Ren, S., Hinzman, A. A., Kang, E. L., Szczesniak, R. D., & Lu, L. J. (2015). Computational and statistical analysis of metabolomics data. Metabolomics, 11(6), 1492–1513. 10.1007/s11306-015-0823-6

Salgado, A. L., Glassmire, A. E., Sedio, B. E., Diaz, R., Stout, M. J., Čuda, J., Pyšek, P., Meyerson, L. A., & Cronin, J. T. (2023). Metabolomic Evenness Underlies Intraspecific Differences Among Lineages of a Wetland Grass. Journal of Chemical Ecology, 49(7), 437–450. 10.1007/s10886-023-01425-2

Samanta, S., Seth, C. S., & Roychoudhury, A. (2024). The molecular paradigm of reactive oxygen species (ROS) and reactive nitrogen species (RNS) with different phytohormone signaling pathways during drought stress in plants. Plant Physiology and Biochemistry, 206, 108259.

Schwachtje, J., Whitcomb, S. J., Firmino, A. A. P., Zuther, E., Hincha, D. K., & Kopka, J. (2019). Induced, Imprinted, and Primed Responses to Changing Environments: Does Metabolism Store and Process Information? Frontiers in Plant Science, *Volume* 10-2019. https://www.frontiersin.org/journals/plant-science/articles/10.3389/fpls.2019.00106

Sharma, A., Das, N., Pandey, P., & Shukla, P. (2025). Plant-microbiome responses under drought stress and their metabolite-mediated interactions towards enhanced crop resilience. Current Plant Biology, 43, 100513. 10.1016/j.cpb.2025.100513

Sharma, M., Kumar, P., Verma, V., Sharma, R., Bhargava, B., & Irfan, M. (2022). Understanding plant stress memory response for abiotic stress resilience: Molecular insights and prospects. Plant Physiology and Biochemistry, 179, 10–24. 10.1016/j.plaphy.2022.03.004

Sun, D., Lei, Z., Carriquí, M., Zhang, Y., Liu, T., Wang, S., Song, K., Zhu, L., Zhang, W., & Zhang, Y. (2025). Reductions in mesophyll conductance under drought stress are influenced by increases in cell wall chelator-soluble pectin content and denser microfibril alignment in cotton. Journal of Experimental Botany, 76(4), 1116–1130.

Szepesi, Á., & Szőllősi, R. (2018). Chapter 17 - Mechanism of Proline Biosynthesis and Role of Proline Metabolism Enzymes Under Environmental Stress in Plants. In P. Ahmad, M. A. Ahanger, V. P. Singh, D. K. Tripathi, P. Alam, & M. N. Alyemeni (Eds.), Plant Metabolites and Regulation Under Environmental Stress (pp. 337–353). Academic Press. 10.1016/B978-0-12-812689-9.00017-0

Tayyab, N., Naz, R., Yasmin, H., Nosheen, A., Keyani, R., Sajjad, M., Hassan, M. N., & Roberts, T. H. (2020). Combined seed and foliar pre-treatments with exogenous methyl jasmonate and salicylic acid mitigate drought-induced stress in maize. PLOS ONE, 15(5), e0232269-. 10.1371/journal.pone.0232269

Tian, H., Bai, J., An, Z., Chen, Y., Zhang, R., He, J., Bi, X., Song, Y., & Abliz, Z. (2013). Plasma metabolome analysis by integrated ionization rapid-resolution liquid chromatography/tandem mass spectrometry. Rapid Communications in Mass Spectrometry, 27(18), 2071–2080. 10.1002/rcm.6666

Tsugawa, H., Cajka, T., Kind, T., Ma, Y., Higgins, B., Ikeda, K., Kanazawa, M., VanderGheynst, J., Fiehn, O., & Arita, M. (2015). MS-DIAL: data-independent MS/MS deconvolution for comprehensive metabolome analysis. Nature Methods, 12(6), 523–526. 10.1038/nmeth.3393

Wasternack, C., & Strnad, M. (2019). Jasmonates are signals in the biosynthesis of secondary metabolites—Pathways, transcription factors and applied aspects—A brief review. New Biotechnology, 48, 1–11.

Wild, J., Kopecký, M., Macek, M., Šanda, M., Jankovec, J., & Haase, T. (2019). Climate at ecologically relevant scales: A new temperature and soil moisture logger for long-term microclimate measurement. Agricultural and Forest Meteorology, 268, 40–47. 10.1016/j.agrformet.2018.12.018

Yu, X., Zhang, W., Zhang, Y., Zhang, X., Lang, D., & Zhang, X. (2019). The roles of methyl jasmonate to stress in plants. Functional Plant Biology, 46(3), 197–212. 10.1071/FP18106

Zeng, G., Gao, F., Xie, R., Lei, B., Wan, Z., Zeng, Q., & Zhang, Z. (2024). The ameliorative effects of exogenous methyl jasmonate on grapevines under drought stress: Reactive oxygen species, carbon and nitrogen metabolism. Scientia Horticulturae, 335, 113354. 10.1016/j.scienta.2024.113354

Zhang, J., Yang, X., Zhang, X., Zhang, L., Zhang, Z., Zhang, Y., & Yang, Q. (2022). Linking environmental signals to plant metabolism: The combination of field trials and environment simulators. Molecular Plant, 15(2), 213–215. 10.1016/j.molp.2021.12.017

Zhou, H., Zhou, G., He, Q., Zhou, L., Ji, Y., & Zhou, M. (2020). Environmental explanation of maize specific leaf area under varying water stress regimes. Environmental and Experimental Botany, 171, 103932.

